# Surviving hypothermia by ferritin-mediated iron detoxification

**DOI:** 10.1101/2021.01.30.428937

**Authors:** Tina Pekec, Jaroslaw Lewandowski, Alicja A. Komur, Daria Sobanska, Yanwu Guo, Karolina Świtońska-Kurkowska, Marcin Frankowski, Maciej Figiel, Rafal Ciosk

## Abstract

How animals rewire cellular programs to survive cold is a fascinating problem with potential biomedical implications, ranging from emergency medicine to space travel. Studying a hibernation-like response in the free-living nematode *Caenorhabditis elegans*, we uncovered a regulatory axis that enhances the natural resistance of nematodes to severe cold. This axis involves conserved transcription factors, DAF-16/FoxO and PQM-1, which jointly promote cold survival by upregulating FTN-1, a protein related to mammalian Fth1/ferritin. Moreover, we show that inducing expression of Fth1 also promotes cold survival of mammalian neurons, a cell type particularly sensitive to deterioration in hypothermia. Our findings in both animals and cells suggest that FTN-1/Fth1 facilitates cold survival by detoxifying ROS-generating iron species. We finally show that mimicking the effects of FTN-1/Fth1 with drugs protects neurons from cold-induced degeneration, opening a potential avenue to improved treatments of hypothermia.

## INTRODUCTION

Cold is a potentially lethal hazard. Nonetheless, hibernation is a widespread phenomenon, used by animals to survive periods of low energy supply associated with cold ^1–4^. Although humans do not hibernate, some primates do so ^5^, hinting that a hibernation-like state might, one day, be induced also in humans, with fascinating medical repercussions ^6, 7^. Nowadays, cooling is used widely in organ preservation for transplantation. Therapeutic hypothermia is also applied, among others, during stroke or trauma, helping preserve functions of key organs, like the brain or heart ^8, 9^. Cellular responses to cold are also of interest for longevity research, as both poikilotherms (animals with fluctuating body temperature, like flies and fish) and homeotherms (like mice) live longer at lower temperatures ^10, 11^. Therefore, understanding the molecular underpinnings of cold resistance has the potential to transform several areas of medicine.

The free-living nematode *C. elegans* populates temperate climates ^12^, indicating that these animals can survive spells of cold. In laboratories, *C. elegans* are typically cultivated between 20-25°C, and a moderate temperature drop slows down, but does not arrest, these animals ^13, 14^. Deep cooling of *C. elegans*, i.e. to near-freezing temperatures, remains less studied. Exposing nematodes to 2-4°C, after transferring them directly from 20-25°C (which we refer to as “cold shock”), results in the death of most animals within one day of rewarming ^15–17^. However, the lethal effects of cold shock can be prevented when animals are first subjected to a transient “cold acclimatization/adaptation” at an intermittent temperature of 10-15°C ^15, 17^. Such cold-adapted nematodes can survive near-freezing temperatures for many days ^15, 17–19^. While in the cold, the nematodes stop aging, suggesting that they enter a hibernation-like state ^17^.

Among factors promoting *C. elegans* survival in near-freezing temperatures, we identified a ribonuclease, REGE-1, homologous to the human Regnase-1/MCPIP1 ^17, 20^. In addition to ensuring cold resistance, REGE-1 promotes the accumulation of body fat, which depends on the degradation of mRNA encoding a conserved transcription factor, ETS-4 ^17^. Interestingly, previous studies showed that the loss of ETS-4 synergizes with the inhibition of insulin signaling in extending lifespan ^21^, and that the inhibition of insulin pathway dramatically enhances cold survival ^15, 19^. Combined, these observations suggested that the cold survival-promoting function of REGE-1 could be related to the inhibition of ETS-4/insulin signaling axis. Here, we validate that hypothesis, dissect the underlying mechanism, and demonstrate that its main objective is the detoxification of harmful iron species. We find that a similar mechanism appears to protect from cold also mammalian cells. By mimicking its effects with drugs, we highlight potential benefits of iron management for treating hypothermia, for which no robust drug treatment currently exists.

## RESULTS

### Inhibition of ETS-4 improves *C. elegans* survival in the cold

Our initial studies of *C. elegans* “hibernation” identified the RNase REGE-1 as a factor promoting cold survival ^17^. Studying REGE-1 in a different physiological context, the regulation of body fat, we showed that a key target of REGE-1 encodes a conserved transcription factor, ETS-4 ^17^. Thus, we asked whether overexpression of ETS-4, taking place in *rege-1(-)* mutants, is also responsible for the cold sensitivity of *rege-1(-)* mutants. We tested that by incubating animals at 4°C, henceforth simply the “cold” (for details on cold survival assay see Fig. 1A). Indeed, we found that *rege-1(-); ets-4(-)* double mutants survived cold much better than *rege-1(-)* single mutants (Fig. 1B). Unexpectedly, we observed that the double mutants survived cold even better than wild type (Fig. 1B). Intrigued, we additionally examined the *ets-4(-)* single mutants and found that they survived cold as well as the double mutants (Fig. 1B). Thus, inhibiting ETS-4 is beneficial for cold survival irrespective of REGE-1. This observation was somewhat surprising as, in wild type, REGE-1 inhibits ETS-4 by degrading its mRNA. However, we observed that, in wild type, both ETS-4 protein and *ets-4* mRNA were more abundant in the cold (Fig. S1A-B). Thus, an incomplete/inefficient degradation of *ets-4* mRNA in the cold could explain the enhanced cold survival of *ets-4(-)* mutants.

**Figure 1.**
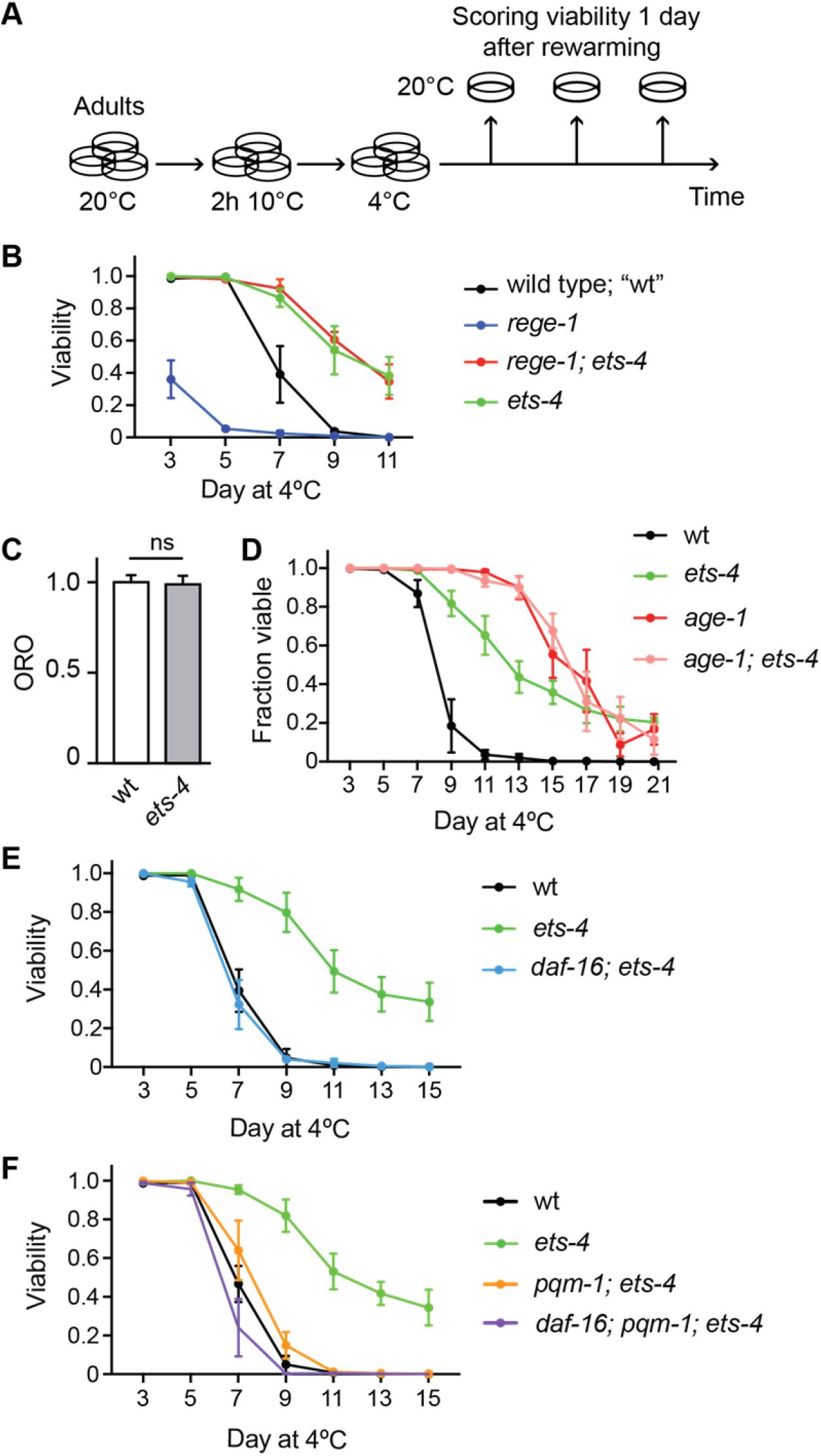
Enhanced cold survival of *ets-4* (-) mutants depends on DAF-16 and PQM-1. **A.** Graphical view of a typical cold-survival experiment, described in detail in the Methods. Shortly, 1 day-old adults, pre-grown at 20°C and distributed between multiple plates, were cold-adapted (for 2 hours at 10°C) and then shifted to 4°C. Every few days, a plate was transferred to 20°C and, after 1 day of recovery at 20°C, the animals were scored for viability. **B.** Cold survival of animals of the indicated genotypes. Mutant alleles throughout the paper are indicated in parentheses and, unless specified otherwise, are loss-of-function alleles. Note that *ets-4(rrr16)* mutants, like *rege-1(rrr13); ets-4(rrr16)* double mutants, survived cold better than wild type (wt). The experiment was performed three times (n= 3); 200-350 animals were scored per time point. Error bars represent standard error of the mean (SEM). **C.** Quantification of body fat, stained with the lipophilic dye oil red O (ORO), in animals of the indicated genotypes. The levels of body fat were similar between wt and *ets-4(rrr16)* mutants. n= 3; 10-15 animals were scored per replicate. Error bars represent SEM. Unpaired two-tailed t-test was used to calculate the p value, “ns” = not significant. **D.** Cold survival of animals of the indicated genotypes (D-F; error bars represent SEM). The *age-1(hx546)* mutants displayed greatly enhanced cold resistance, and combining *age-1(hx546)* and *ets-4(rrr16)* mutations did not provide animals with additional resistance. n= 4; 350-500 animals were scored per time point. **E.** Combining *daf-16(mu86)* and *ets-4(rrr16)* mutations abolished the enhanced cold survival of *ets-4(-)* mutants, reverting it to wild-type values. n= 4; 350-500 animals were scored per time point. **F.** Combining *pqm-1(ok485)* and *ets-4(rrr16)* mutations abolished the enhanced cold survival of *ets-4(-)* mutants, reverting it to wild-type values. Also note that the triple *daf-16(mu86); pqm-1(ok485); ets-4(rrr16)* mutants survived cold essentially like wild type, indicating that DAF-16 and PQM-1 promote cold survival in *ets-4(-)*, but not wild-type animals. n= 4; 450-650 animals were scored per time point.

Because many hibernators burn fat to fuel survival in the cold, the ETS-4-mediated fat loss ^17^ and cold sensitivity (reported here), observed in *rege-1(-)* mutants, could be connected. However, we found that inhibiting ETS-4 restores body fat of *rege-1(-)* mutants to only wild-type levels ^17^, and yet the *rege-1(-); ets-4(-)* double mutants are more resistant to cold than wild type (Fig. 1B). We additionally examined the fat content of *ets-4(-)* single mutants and found that it was indistinguishable from wild-type (Fig. 1C). Thus, ETS-4 appears to impact body fat and cold resistance via separate mechanisms.

### The enhanced cold survival requires both DAF-16 and PQM-1

ETS-4 was previously described to synergize with the insulin/IGF-1 signaling pathway in limiting the nematode lifespan ^21^. Moreover, the lifespan extension seen in *ets-4(-)* mutants, as is the case with insulin pathway mutants, depends on the transcription factor DAF-16/FOXO ^21, 22^. These and additional reports, that insulin pathway mutants display cold resistance depending on DAF-16 ^15, 19^, prompted us to examine the genetic relationship between *ets-4(-)* and insulin pathway mutants in the context of cold resistance. Firstly, using a loss-of-function allele of the insulin-like receptor, *daf-2(e1370)* ^23^, we confirmed that inhibiting insulin signaling improves cold survival (Fig. S1C). Combining this *daf-2* mutation with *ets-4(-),* we observed that the double mutants survived cold slightly better than the *daf-2* single mutant (Fig. S1C). However, the difference between *daf-2(e1370); ets-4(-)* and *daf-2(e1370)* mutants was smaller than the difference between *ets-4(-)* mutant and wild type, suggesting a partial overlap between mechanisms activated upon the inhibition of DAF-2 or ETS-4.

The partial overlap suggests that ETS-4 may affect insulin signaling “downstream” from the DAF-2 receptor. The main components of the *C. elegans* insulin pathway include the phosphoinositide 3-kinase AGE-1/PI3K, which is why we also tested the genetic relationship between *age-1* and *ets-4* mutations. Using the *age-1(hx546)* allele, carrying a point mutation reducing the AGE-1 activity ^24^, we confirmed that also the *age-1(-)* mutants survive cold better than the wild type, and that their improved survival depends on the transcription factor DAF-16/FOXO (Fig. S1D). Then, we examined the epistatic relationship between *age-1(hx546)* and *ets-4(-)* mutants. While the *age-1(hx546)* single mutants survived cold, expectedly, much better than wild type (Fig. 1D), we observed no additional benefit of combining *age-1(hx546)* and *ets-4(-)* mutations (Figs. 1D and S1D). These observations suggest that AGE-1 could act in the same pathway as ETS-4, or alternatively converge on the same downstream effector(s). Thus, we also examined whether the enhanced cold survival of *ets-4(-)* mutants depends on DAF-16. Indeed, we found that removing DAF-16 completely suppressed the enhanced cold survival of *ets-4(-)* mutants (Fig. 1E). Reconciling all observations, we hypothesize that, in wild type, signals generated upon DAF-2 or ETS-4 activation converge on AGE-1, thus inhibiting DAF-16 and limiting cold resistance. Conversely, upon the inactivation of DAF-2 or ETS-4, DAF-16 activation results in improved cold resistance.

Recently, another transcription factor, PQM-1, was shown to complement DAF-16 in promoting the lifespan in DAF-2 deficient animals ^25^. In the intestinal cells, PQM-1 and DAF-16 nuclear occupancy has been shown to be mutually exclusive, and they appear to regulate largely separate sets of target genes ^25^. Additionally, these transcription factors can play opposing roles; for example, while the formation of an alternative “dauer” larval stage (deployed to survive adverse environmental conditions) depends on DAF-16 ^22^, PQM-1 facilitates the recovery from the dauer arrest ^25^. Yet there is also some evidence supporting synergistic roles of DAF-16 and PQM-1. For example, both the class I and II genes (see the introduction) are down-regulated in *pqm-1* mutants ^25^, and DAF-16 and PQM-1 both contribute to the lifespan extension of *daf-2* or mitochondrial mutants ^25, 26^. Thus, at least in certain circumstances, DAF-16 and PQM-1 may collaborate. Therefore, we tested whether the loss of PQM-1 had a similar effect on the cold survival of *ets-4(-)* mutants as the loss of DAF-16, and observed that, indeed, removing PQM-1 suppressed the enhanced cold survival of *ets-4(-)* mutants (Fig. 1F). Importantly, in otherwise wild-type background, we observed no apparent effects on cold survival in either *pqm-1(-)* or *daf-16(-)* single mutants, nor in the *pqm-1(-); daf-16(-)* double mutants (Fig. S1E). Even the survival of *daf-16(-); pqm-1(-); ets-4(-)* triple mutants was indistinguishable from wild-type nematodes (Fig. 1F). Together, these observations argue for a specific, joint role for DAF-16 and PQM-1 in cold survival, under conditions that favor their activation, such as upon ETS-4 inactivation.

### DAF-16 and PQM-1 are enriched in the gut nuclei in the cold

Presumably, DAF-16 and PQM-1 facilitate cold survival by inducing transcription of specific genes. Under normal growth conditions, DAF-16 remains inactive in the cytoplasm. However, when insulin signaling is inhibited, DAF-16 moves to the nucleus to activate target genes. Based on the genetic analysis above, we suspected the nuclear accumulation of DAF-16 in *ets-4(-)* mutants. To examine that, we attached (by CRISPR/Cas9 editing) a GFP-FLAG tag to the C-terminal end of the endogenous *daf-16* ORF (see Methods). Examining the distribution of GFP-tagged DAF-16 (DAF-16::GFP), we observed little nuclear signal at 20°C, possibly with a minimal increase in the absence of ETS-4. After one or three days at 4°C, however, we observed a significant increase in the nuclear DAF-16::GFP (Fig. 2A and C); that increase appeared to be posttranscriptional, as *daf-16* mRNA levels remained constant between 20°C and 4°C (Fig. S2A). Although the nuclear DAF-16::GFP signal appeared slightly stronger in *ets-4(-)* mutants at day one in the cold, that was no longer true at day 3 (Fig. 2A and C). Thus, although the nuclear enrichment of DAF-16 is consistent with its ability to potentiate cold resistance, that enrichment is, apparently, insufficient, as it only enhances cold survival in *ets-4(-)* mutants but not wild type.

**Figure 2.**
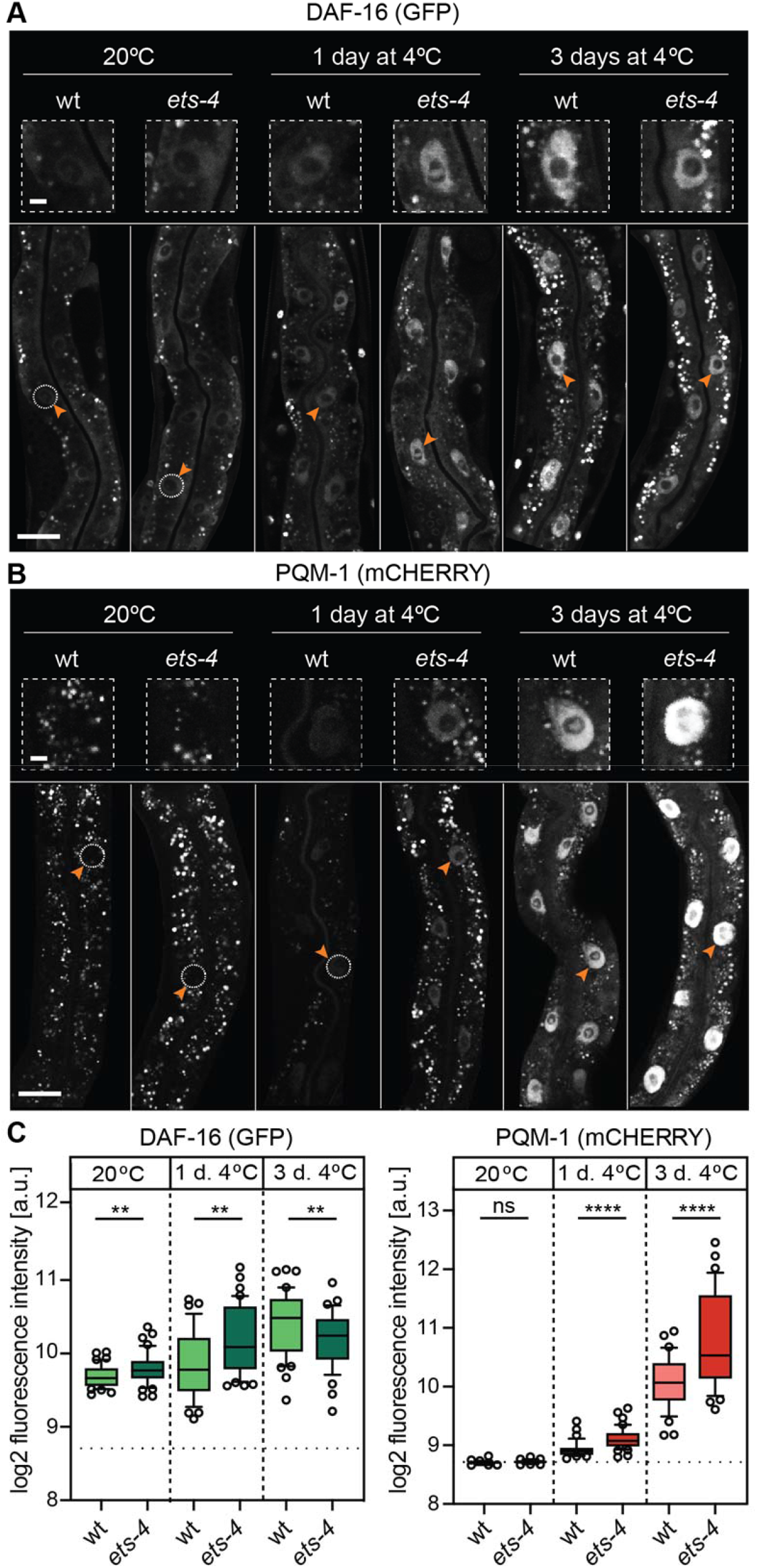
DAF-16 and PQM-1 are enriched in the nuclei in the cold. **A.** Micrographs showing representative confocal images of GFP fluorescence, reflecting the endogenously tagged DAF-16, allele *daf-16(syb707)*, from wt or *ets-4(rrr16)* mutants. The animals were sampled at the indicated times and temperatures, according to 1A. The corresponding quantifications are in C. Arrowheads point to representative gut nuclei (demarcated with dashed circles when displaying little or no fluorescence), which are enlarged in the insets above. Size bars, here and in B: 5 μm (small magnification) and 25 μm (large magnification). **B.** Representative confocal images of mCherry fluorescence, reflecting the endogenously tagged PQM-1, allele *pqm-1(syb432)*, from wt or *ets-4(rrr16)* mutants. The animals were sampled as above. The corresponding quantifications are in C. Arrowheads point to representative gut nuclei, enlarged in the insets above. **C.** Quantifications of the nuclear fluorescence, corresponding to A (left) and B (right). Each data point represents log_2_-transformed mean nuclear intensity per animal. Dotted line represents the average background within each experiment. Left: n= 3; 10 to 15 animals were scored per replicate. Error bars represent 10^th^ to 90^th^ percentile. Unpaired two-tailed t-test was used to calculate the p value. ** indicates *p* < 0.01. Right: n= 3; 10 to 15 animals were scored per replicate. “ns” = not significant; **** indicates *p* < 0.0001.

Thus, we performed the same analysis on PQM-1, fusing (by CRISPR/Cas9 editing) an mCHERRY-MYC tag to the C-terminal end of the endogenous *pqm-1* ORF (see Methods). We detected little, if any, nuclear PQM-1::mCHERRY at 20°C, in either wild-type adult nematodes or *ets-4(-)* mutants (Fig. 2B-C), agreeing with the previously reported expression patterns ^25, 27–29^. By contrast, after one day at 4°C, we began detecting the nuclear PQM-1::mCHERRY signal in wild-type nematodes, and a slightly stronger signal in the nuclei of *ets-4(-)* mutants (Fig. 2B-C); this increase may be transcriptional, as *pqm-1* mRNA levels were higher at 4°C than 20°C (Fig. S2B). After three days at 4°C, the PQM-1::mCHERRY nuclear signal increased even further (Fig. 2B-C) and, at this point, *ets-4(-)* mutants displayed significantly higher signal than wild type (Fig. 2B-C). Thus, in contrast to the standard cultivation temperature, where DAF-16 and PQM-1 localize to the nucleus in a mutually exclusive manner ^25^, DAF-16 and PQM-1 coexist in the nucleus in the cold. Intriguingly, DAF-16 and PQM-1 induce FTN-1 in *ets-4* mutants but not wild type. Thus, one possibility is that the additional accumulation of PQM-1 (observed in the absence of ETS-4) is required to reach a threshold for DAF-16 activation. Among other possibilities, DAF-16 activation could involve additional factors normally inhibited by ETS-4.

### Identification of a PQM-1 and DAF-16 coregulated gene promoting cold survival

The above observations are compatible with a scenario where, upon ETS-4 inactivation, DAF-16 and PQM-1 coregulate transcription of cold survival-promoting gene(s). To test this hypothesis, we undertook a functional genomic approach. First, we compared gene expression (by RNAseq) between *ets-4(-)* and wild-type animals incubated at 4°C. Then, by comparing *pqm-1(-); ets-4(-)*, or *daf-16(-); ets-4(-)* double mutants to the *ets-4(-)* single mutant, we identified genes, whose expression in the *ets-4(-)* mutant depends on PQM-1 and/or DAF-16. To illustrate this, we prepared an integrative heat-map, using all 4°C samples with replicates. Focusing on changes between the strains, we observed 3 distinct clusters (Fig. S3A). Cluster 1 (red) includes genes upregulated in the cold in *ets-4(-)* mutants (compared to wild type), which either do not change or go down, upon the additional inhibition of *daf-16* or *pqm-1*. Cluster 2 (green) includes genes upregulated across all conditions. Finally, the smallest cluster 3 (blue), includes genes downregulated in *ets-4(-)* mutants, which either do not change or go up, upon the additional inhibition of *daf-16* or *pqm-1*. With this analysis, we observed that many changes in gene expression upon the loss of ETS-4, were reverted upon the additional loss of either DAF-16 or PQM-1, supporting a functional relationship between DAF-16 and PQM-1. Taking advantage of the ENCODE database, which reports genome-wide chromatin association of many transcription factors ^30^, we examined the potential binding of DAF-16 and PQM-1 around the transcription start sites (TSS) of genes in each cluster of the heat map. Even though the ENCODE data comes from experiments performed at standard growth conditions, we decided to use it as an approximation, and observed that genes, whose expression in *ets-4(-)* mutants depends on DAF-16 or PQM-1 (i.e. genes in clusters 1 and 3), appear to be enriched for TSS-proximal binding sites for both transcription factors (Fig. S3A). The same enrichment was not seen for the cluster 2 genes, whose expression is apparently unrelated to DAF-16 or PQM-1 (Fig. S3A). The possible connection between clusters 1 and 3, and the association with DAF-16 or PQM-1, was statistically significant for PQM-1 but not DAF-16 (Fig. S3A). Nevertheless, by analyzing transcription factor binding motifs enriched within each gene cluster, we observed a DAF-16-like motif enriched within cluster 1 genes (Fig. S3B), which made us focus on those genes.

To identify candidate genes, whose DAF-16 and PQM-1 dependent activation promotes cold survival, we first selected genes upregulated (in both biological replicates), at least two-fold, in *ets-4(-)* mutants compared to wild type (after one day at 4°C). Second, we intersected these genes with those whose promoters associate with either DAF-1 or PQM-1, according to the confident binding sites from Tepper et al. This analysis yielded seven genes that were reproducibly upregulated in *ets-4(-)* mutants, and whose promoters may associate with both PQM-1 and DAF-16 (Fig. 3A). If these genes were relevant for the enhanced cold survival, their inhibition would be expected to impede cold survival of *ets-4(-)* mutants. Testing this, we observed that RNAi-mediated depletion of one candidate, *ftn-1* (encoding a nematode ferritin), reproducibly compromised cold survival of *ets-4(-)* mutants (Fig. 3B; note that the RNAi construct is predicted to target also *ftn-2*, which is highly similar *to ftn-1*, see below).

**Figure 3.**
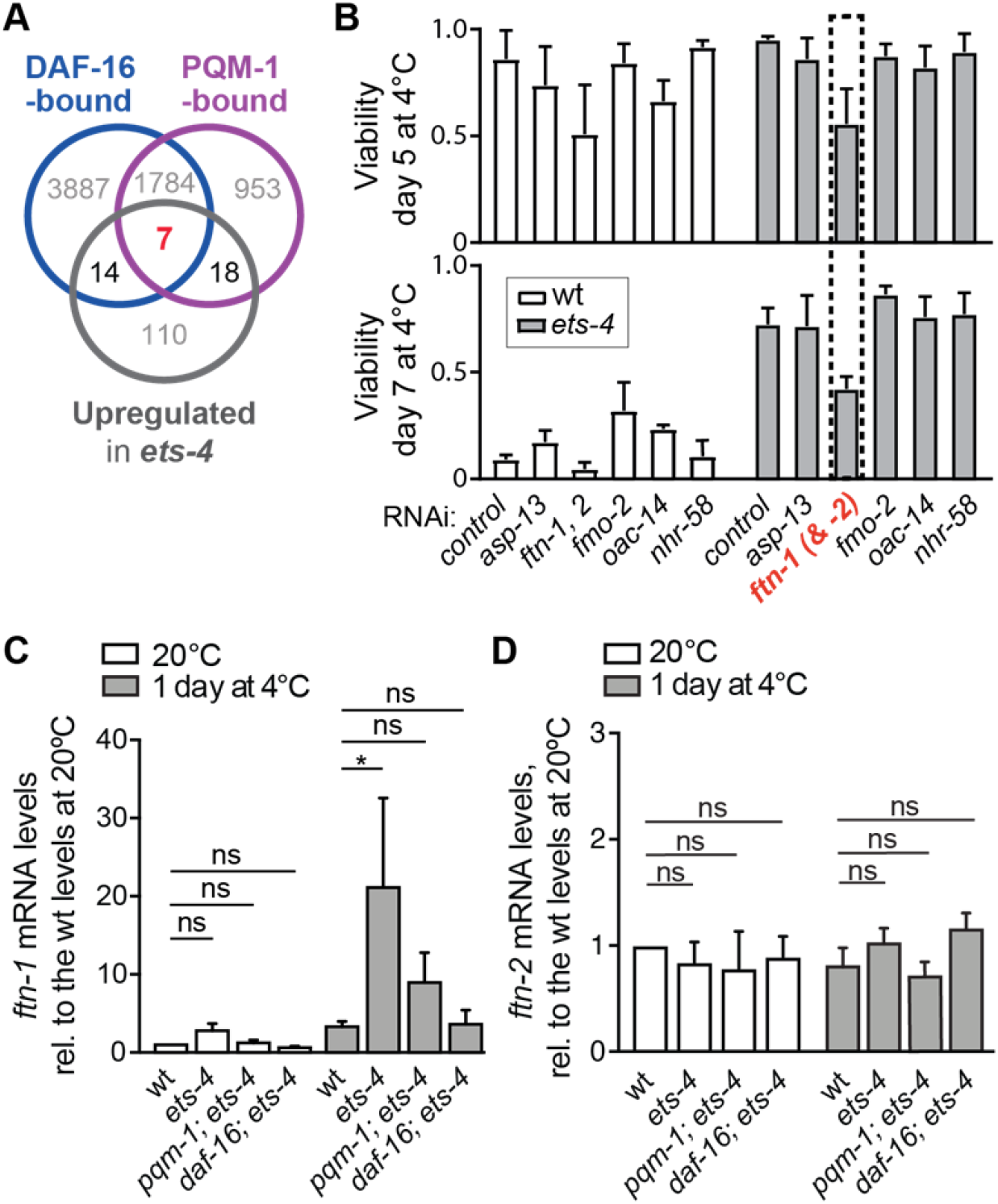
DAF-16 and PQM-1 coregulate expression of *ftn-1/ferritin*. **A.** Diagram comparing relations between three sets of genes: Grey circle: genes upregulated more than 2-fold in *ets-4(rrr16)* mutants compared to wt (at day 1 at 4°C and in two replicates, see Fig. S3A). Blue circle: genes whose promoters are bound by DAF-16 (according to Tepper et al.). Magenta: genes whose promoters are bound by PQM-1 (according to Tepper et al.). Note that seven genes (*asp-13*, *cpt-4*, *fmo-2*, *ftn-1*, *nhr-58*, *oac-14*, and *pals-37*), whose promoters are bound (at normal temperature) by both DAF-16 and PQM-1, were reproductively upregulated in the absence of ETS-4. **B.** The candidates from A were RNAi-depleted, from either wt or *ets-4(rrr16)* animals, and those animals were tested for cold resistance (according to 1A) at the indicated times. Note that depleting *ftn-1* (and *ftn-2*, as RNAi is predicted to target both homologs) significantly reduced cold survival of *ets-4(rrr16)* animals (stippled box). Error bars represent SEM. n= 3; 200-350 animals were scored per time point. **C.** Shown are *ftn-1* mRNA levels, measured by RT-qPCR, in animals of the indicated genotypes. Strains used: wt, *ets-4(rrr16)*, *pqm-1(ok485); ets-4(rrr16)*, and *daf-16(mu86); ets-4(rrr16).* The animals were sampled at 20°C, before cold adaptation, and after one day at 4°C, according to 1A. The mRNA levels were normalized to the levels of *act-1* (actin) mRNA. At each temperature, the values were then normalized to those from the wild type at 20°C. n= 5; error bars represent SEM. P values were calculated using 2-way ANOVA for multiple comparisons. “ns” = not significant; * indicates p > 0.05. **D.** The animals were collected as in C. The levels of *ftn-2* mRNA, measured by RT-qPCR, were normalized to *act-1* mRNA, and are shown relative to the *ftn-2* mRNA level in wt at 20°C. Wt and *ets-4(-)* animals were collected at 20°C and after one day at 4°C (n= 3). P values were calculated using 2-way ANOVA. Error bars represent SEM. “ns” = not significant.

### FTN-1/Ferritin promotes cold survival

Iron is an essential but also a potentially harmful element, whose cellular levels are tightly regulated by various molecular mechanisms, which are largely conserved between nematodes and humans ^31, 32^. Among critical iron regulators are iron-binding proteins called ferritins. In *C. elegans*, ferritin is encoded by two genes, *ftn-1* and *ftn-2*. Under standard growth conditions, *ftn-2* mRNA is much more abundant than *ftn-1* ^33–35^. While *ftn-1* was previously shown to be upregulated upon DAF-2 inactivation, in a DAF-16 dependent manner, *ftn-2* is not considered a DAF-16 target ^36^. Also, in agreement with their differential regulation in innate immunity, the expression of *ftn-1*, but not *ftn-2*, was reported to depend on PQM-1 ^28^. Confirming our RNA profiling data by RT-qPCR, and consistent with the regulation by both DAF-16 and PQM-1, we observed a DAF-16 and PQM-1 dependent increase in the levels of *ftn-1* mRNA in cold-treated *ets-4(-)* mutants (Fig. 3C-D). Because the RNAi construct targets both *ftn-1* and *ftn-2*, we also examined FTN-1 function using an existing loss-of-function allele, *ftn-1(ok3625)* ^37, 38^. Since DAF-16 and PQM-1 are important for cold survival in *ets-4(-)*, but not wild-type animals, their relevant target may be expected to display a similar behavior. Indeed, the cold survival of *ftn-1(-)* single mutants was indistinguishable from wild type (Fig. 4A), but the *ftn-1* inactivation completely abolished the enhanced cold survival of *ets-4(-)* mutants (Fig. 4A). Importantly, FTN-1 is expressed in the intestine ^34^, i.e. the tissue where ETS-4, DAF-16 and PQM-1 are all expressed.

**Figure 4.**
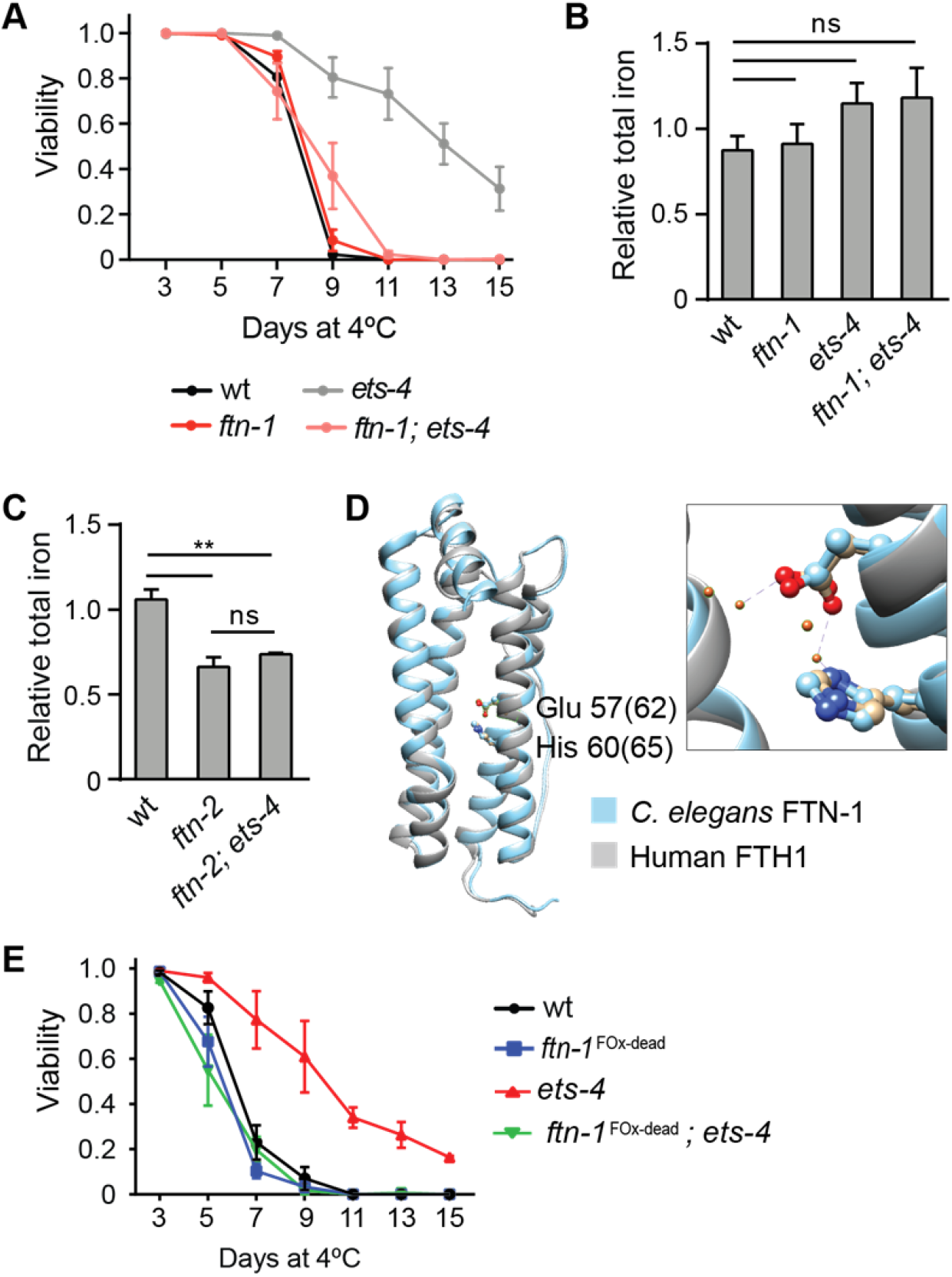
FTN-1 promotes cold survival via its ferroxidase activity. **A.** Viability of animals, of the indicated genotypes, subjected to cold as explained in 1A. Error bars represent SEM. n= 3; 250-400 animals were scored per time point. Note that combining the *ftn-1(ok3625)* mutation with *ets-4(rrr16)* reverted the enhanced cold survival of *ets-4(-)* mutants to wild-type values, similar to the double *daf-16(-); ets-4(-)* or *pqm-1(-); ets-4(-)* mutants in 1E-F. Also, like *daf-16(-)* and *pqm-1(-)* single mutants in S1E, *ftn-1(ok3625)* single mutants survived cold as well as wt. **B.** Total iron levels in *C. elegans* extracts, measured by ICP-MS. Wild type, *ets-4(rrr16), ftn-1(ok3625)*, and *ftn-1(ok3625); ets-4(rrr16)*, 1 day-old adults were subjected to cold for 3 days. The values were normalized to those of the wild type. Unpaired two-tailed t-test was used to calculate the p value, “ns” = not significant. Error bars represent SEM, n= 3. Note that inactivating FTN-1 had no impact on the total iron levels. **C.** Comparing total iron levels, measured by ICP-MS, in *C. elegans* extracts derived from wild-type, *ftn-2*(*ok404*), and *ftn-2*(*ok404*); *ets-4*(*rrr16*), 1 day-old adults, subjected to cold for 3 days. The values were normalized to those of the wild type. Unpaired two-tailed t-test was used to calculate the p value, “ns” = not significant, ** p value < 0.01. Error bars represent SEM, n= 3. Note that inactivating FTN-2 decreased the levels of total iron, also in *ets-4(-)* animals. **D.** Structural alignment of *H. sapiens* ferritin heavy chain 1 (FTH1 – colored in grey, PDB code 4OYN) and *C. elegans* FTN-1 (colored in light blue), using the Phyre2 tool ^64^. Amino acids critical for the ferroxidase activity are shown as balls and sticks. The colors indicate: carbon atoms in FTH1 (light brown) or FTN-1 (light blue), oxygen atom of glutamic acid (red), and nitrogen atom of histidine (dark blue). The magnification shows the ferroxidase active site, with the iron atoms shown as dark orange balls and coordination bonds as dotted lines. **E.** Cold survival of animals of the indicated genotypes subjected to cold, as in 1A. “*ftn-1*^FOx-dead^” indicates the *ftn-1*(*syb2550*) allele, which encodes FTN-1 variant with point mutations (E57K and H60G, where the first methionine is counted as 0) abolishing its ferroxidase activity. Note that combining *ets-4(rrr16)* and *ftn-1*(*syb2550*) mutations completely abolished the enhanced cold resistance of *ets-4(-)* single mutant. Error bars represent SEM. n= 3; 261-396 animals were scored per time point.

If the cold survival-enhancing role of FTN-1 is related, as expected, to its function in iron regulation, excess iron may be expected to impede cold survival. We tested that by supplementing culture plates with ferric ammonium citrate (FAC). Although we do not know how much extra iron is being absorbed by FAC-treated animals, we observed a dose-dependent impediment of cold survival (Fig. S4A). Importantly, although higher FAC levels reduced the survival of *ets-4(-)* mutants, their survival was still greater than of the corresponding (i.e. FAC-treated) wild type (Fig. S4B). The better survival of FAC-treated *ets-4(-)* mutants still depended on FTN-1, as the *ftn-1(-); ets-4(-)* double mutants responded to excess iron like the corresponding wild type (Fig. S4B). Combined, our data suggest that FTN-1, when expressed in cold-treated *ets-4(-)* animals, facilitates survival, and its beneficial effect involves some form of iron regulation.

### FTN-1 promotes cold survival through its ferroxidase activity

Mammalian ferritin consists of multiple heavy and light subunits (FTH and FTL) that form nanocages storing thousands of iron atoms ^39^. The *C. elegans* ferritins, FTN-1 and −2, are more similar to FTH ^33^. To test iron sequestration by FTN-1, we used size exclusion chromatography-inductively coupled plasma-mass spectrometry (SEC-ICP-MS; ^38^). We found that, in cold-treated animals, the levels of total iron were independent of FTN-1 (Figs. 4B, S4C, E) but, as reported at standard temperature ^38^, depended on FTN-2 (Figs. 4C, S4D, E). Thus, FTN-1 appears to contribute little, if at all, to the pool of stored iron.

Iron is present in cells in both oxidized Fe^3+^/ferric(III) and reduced Fe^2+^/ferrous(II) forms. Excess of Fe^2+^ is potentially harmful, because, in the so-called Fenton reaction, it catalyzes the formation of reactive oxygen species, ROS ^40, 41^. Notably, both FTN-1 and −2 contain predicted ferroxidase active sites, which in homologous proteins mediate the Fe^2+^-to-Fe^3+^ conversion (Figs. 4D and S4F). Accordingly, the *ftn-2; ftn-1* double mutants display an elevated ratio of Fe2+/Fe3+ during aging ^38^. Although the individual impact of FTN-1 on the Fe2+/Fe3+ balance was not examined, the overexpression of *ftn-1* was reported to have antioxidant effects ^42^. To test whether the ferroxidase activity of FTN-1 promotes cold survival, we modified the endogenous *ftn-1* locus (by CRISPR/Cas9 editing), so that it produces a ferroxidase-dead FTN-1. Crucially, inactivation of the FTN-1 ferroxidase activity completely abolished the enhanced cold survival of *ets-4(-)* animals (Fig. 4E). Thus, FTN-1 appears to facilitate cold survival through iron detoxification, rather than sequestration.

### Overproduction of FTN-1 is sufficient for the enhanced cold survival

Thus far, we have shown that FTN-1, when expressed in the absence of ETS-4, gives worms advantage in surviving cold. To test whether FTN-1 may do that in otherwise wild-type background, we created (using Mos1-mediated Single Copy Insertion, MosSCI; ^43^) strains overexpressing *ftn-1* from two different, robust promoters, *dpy-30* and *vit-5*. Importantly, we found that both strains survived cold much better than wild type (Fig. 5A; for the levels of *ftn-1* mRNA overproduced from the *vit-5* promoter see Fig. S5A). By SEC-ICP-MS, we observed no changes in the levels of ferritin-associated iron in the overexpressing strains (Figs. 5B and S5B, C), consistent with FTN-1 functioning in iron detoxification and not sequestration.

**Figure 5.**
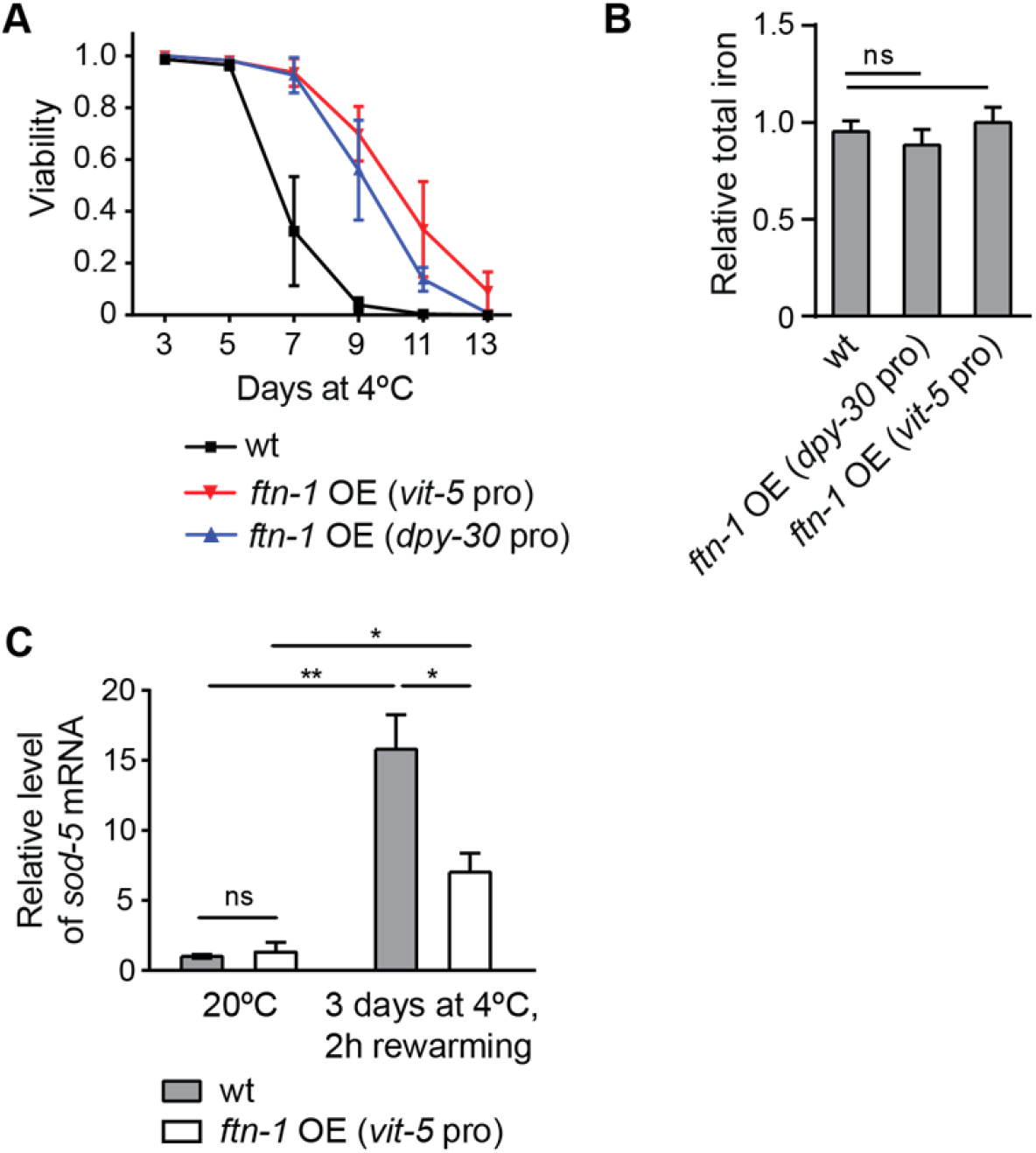
FTN-1 overexpression is sufficient for enhanced cold survival. **A.** Survival of animals of the indicated genotypes subjected to cold, as in 1A. The *ftn-1* overexpressing lines (*sybSi67* and *sybSi72*) are marked as *ftn-*1 OE (*dpy-30* pro) and *ftn-*1 OE (*vit-5* pro), respectively. Note that both *ftn-1*-overexpressing strains survived cold better than wt; *ftn-1* overexpression from the *vit-5* promoter was slightly more beneficial, which is why we chose this strain for additional experiments. Error bars represent SEM, n= 3. 232-307 animals were scored per time point. **B.** Total iron levels, measured by ICP-MS, in *C. elegans* extracts derived from wild type, or *ftn-1* overexpressing, 1 day-old adults, grown at 20°C. The values were normalized to wild type. Unpaired two-tailed t-test was used to calculate the p value, “ns” = not significant. Error bars indicate SEM, n= 3. **C.** Relative levels of *sod-5* mRNA, measured (by RT-qPCR) in animals of the indicated genotypes, and at the time and temperature. Note that the cold-treated animals were analyzed at around 2 hours into rewarming at 20°C. mRNA levels were normalized to *act-1* mRNA. At each time point, the values were then normalized to the wt at 20°C. Error bars represent SEM. n= 3; p values were calculated using an unpaired Student t-test. * indicates p < 0.05; ** p < 0.01; “ns” = not significant.

The ferroxidase activity of FTN-1 is expected to lower the levels of ROS-generating Fe(II), implying that cold-treated nematodes experience increased levels of ROS. We found that reagents typically used for ROS detection (like CM-H_2_DCFDA) are toxic to cold-treated animals. Thus, we sought a factor whose induction could be used as a proxy for ROS detection. Specific enzymes, called superoxide dismutases (SODs), function at the front line of cellular defense against ROS ^44^. There are five SODs in *C. elegans* and, examining their expression in cold-treated animals, we noticed a consistent increase in the *sod-5* mRNA. To understand the dynamics of *sod-5* activation, we examined the expression of GFP-tagged SOD-5 ^45^; the fusion protein is expressed mainly in neurons. Following SOD-5::GFP signal in live animals, we observed a strong, but transient increase of SOD-5::GFP during rewarming (Fig. S5D; note the elevated signal around 2 h into rewarming). Focusing thus on this time point, we tested whether the overexpression of *ftn-1* impacts *sod-5* activation. Indeed, we observed that the levels of *sod-5* mRNA were significantly lower in the *ftn-1* overexpressing strain than wild type (Fig. 5C). All observations combined, a picture emerges where FTN-1, through its iron(II)-detoxifying activity, protects animals from the cold by reducing the levels of Fe(II)-catalyzed ROS. According to this model, animals subjected to cold experience an increase in Fe(II) iron. Detection of specific iron forms is not trivial, and our attempts to detect specifically Fe(II) in *C. elegans* were unsuccessful. Thus, assuming some level of conservation in cellular responses to cold, we decided to investigate that in mammalian cells, where Fe(II) detection is more robust and potential findings more appllicable to human hypothermia.

### Iron management plays a key role in neuronal resistance to cold

Since the main clinical benefit of deep cooling is the preservation of neuronal functions, we decided to examine Fe(II) in neurons, where our observations may be of clinical relevance. For convenience, we chose to study murine neurons. To generate them, we differentiated primary neuronal stem cells, collected from early mouse embryos, into noradrenergic-like neurons (henceforth “neurons”), which affect numerous physiological functions, generally preparing the body for action. To examine their cold resistance, neurons (cultivated at the physiological temperature of 37°C) were shifted to 10°C for 4 hours and then returned to 37°C. Their viability was examined after rewarming for 24 hours (see Methods for details). First, we observed that cooling induced cell death in a large fraction of neurons (Fig. 6A). Interestingly, neuronal death was associated with rewarming (Fig. S6A), which is somewhat reminiscent of reperfusion injury, arguing that not the cold *per se*, but rather the burden associated with restoring cellular functions during rewarming, is the critical challenge facing cold-treated neurons.

**Figure 6.**
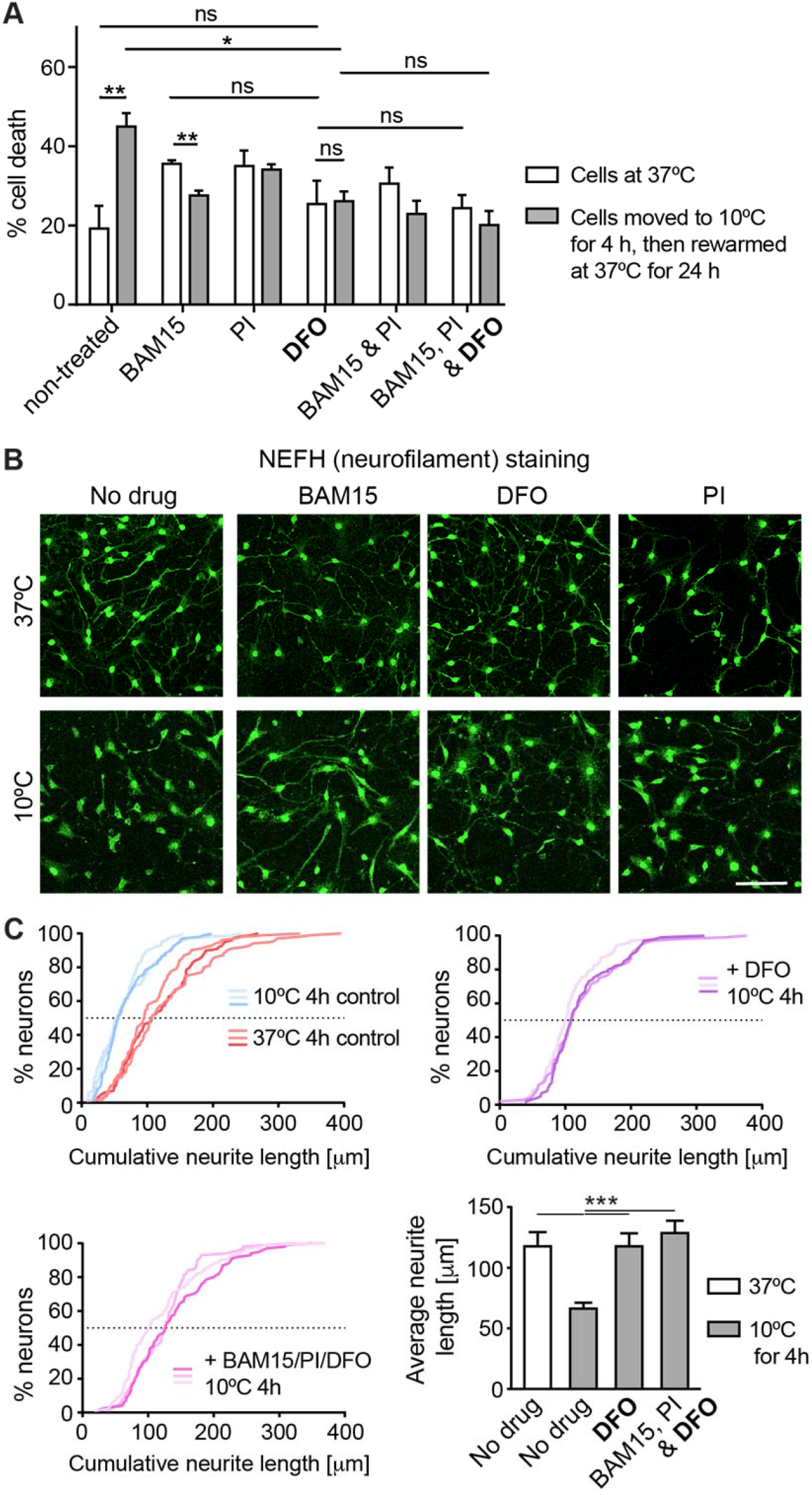
Reducing free iron protects murine neurons from cold-induced degeneration. **A.** Viability of murine neurons, subjected to cold and the indicated drugs, was examined by staining with propidium iodide (see Methods for details). “BAM15” is a mitochondrial uncoupling drug, “PI” a cocktail of protease inhibitors, and “DFO” deferoxamine, an iron chelator. Error bars represent SEM. n= 3 experiments; p values, between cold-treated and control (37°C) samples, were calculated with Student’s t-test, while the ANOVA plus Tukey post hoc test was used to compare samples subjected to different drugs. * indicates p < 0.05; ** p < 0.01; and “ns” not significant. Note that, in contrast to non-treated cells incubated at 10°C, treating cells with DFO prevented cell death to a similar extent as the treatment with BAM15, PI, or the combination of drugs. **B.** Representative confocal images of murine neurons, subjected to cold and the indicated drugs, and immunostained for NEFH to visualize neurites immediately after cold treatment. Scale bar: 40 μm. Note that the neurites, which in control neurons degenerated upon cold exposure (the bottom left panel), were stabilized by adding DFO, similar to BAM15 or PI. **C.** Quantifications of neurite lengths, visualized by NEFH labeling, corresponding to B. The cumulative plots (see Methods) compared neurite lengths in cells treated as indicated. Each curve corresponds to one experimental replicate. The bar graph (bottom right) compares average neurite lengths. While cold treatment led to the shortening of neurites, treating cells with DFO alone stabilized the neurites equally well, as when combined with BAM15 and PI. Error bars represent SD. *** indicates p < 0.001.

A recent study compared cold survival of neurons derived from either hibernating or non-hibernating mammals, and reported that “hibernating” neurons survive cold much better than “non-hibernating” ones ^46^. Thus, hibernating neurons appear to possess intrinsic mechanisms enhancing cold resistance. Remarkably, treating non-hibernating neurons with certain drugs was shown to compensate, at least partly, for their lower cold resistance. Although house mice, upon starvation, are capable of daily torpor ^47^, they can be considered as non-hibernators in a classical sense. Correspondingly, we observed that treating murine neurons with either BAM15 or PI (drugs previously used by Ou et al.) increased their cold survival (Fig. 6A). Assuming that, like for nematodes, iron management is crucial for the survival of cold-treated neurons, we treated neurons with the iron-chelating drug deferoxamine, DFO (expected to lower cellular levels of free iron). Indeed, DFO treatment protected neurons from cold-related death to the same extent as BAM15, PI, or drug combinations (Fig. 6A).

In contrast to hibernating neurons, the non-hibernating neurons display a striking deterioration of neuronal processes/neurites, which is counteracted by BAM15 and/or PI treatment ^46^. By staining neurons against NEFH (neurofilament protein heavy polypeptide; a neuron-specific component of intermediate filaments), we observed that also DFO had a strong stabilizing effect on cold-treated neurites (Fig. 6B-C). This protection appeared to be long-lasting, as the neurites were still evident at 24 h into rewarming (Fig. S6B-C).

### Overproduction of FTH1 improves cold survival of mammalian neurons

By lowering the pool of free iron, DFO could, indirectly, reduce the levels of Fe(II). Nonetheless, to monitor ferrous iron directly, we employed a fluorescent probe, FeRhoNox-1, which specifically detects Fe(II). Strikingly, we observed a strong, though transient, increase of Fe(II) during rewarming (Fig. S7A-B), which was prevented by DFO treatment (Fig. 7A). Since ferrous iron catalyzes the formation of reactive oxygen species (ROS), we also measured ROS levels, using CellROX-green. We observed a strong increase of ROS during rewarming, at the time coinciding with the Fe(II) peak. Importantly, that increase was counteracted by DFO treatment, as expected (Fig. 7B).

**Figure 7.**
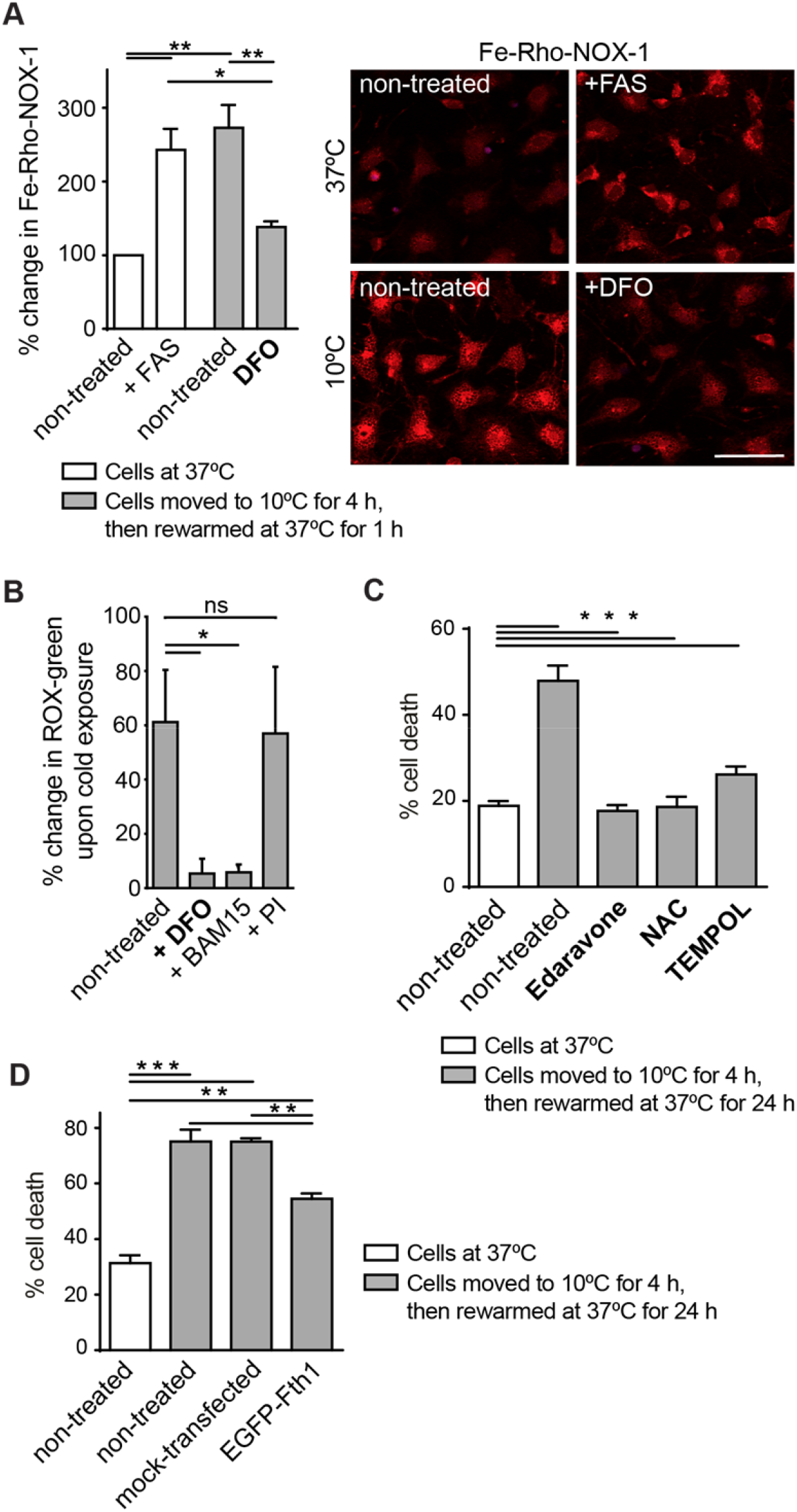
Lowering iron(II) or ROS, or FTH1 overexpression, all enhance neuronal cold resistance. **A.** Examining iron(II) levels, using the fluorescent FeRhoNox-1 probe (see Methods), in neurons subjected to cold and/or the indicated drugs. Left: quantifications of FeRhoNox-1 fluorescence, relative to non-treated cells incubated at 37°C. “FAS” indicates ammonium ferrous sulfate, here a source of additional iron(II). n= 3; p values were calculated by ANOVA plus post hoc Tukey test. ** indicates p < 0.01; and * p < 0.05. Error bars represent SEM. Representative images are on the right. Scale bar: 50 μm. Note that the exposure to cold resulted in higher levels of reactive iron, and that this accumulation was prevented by DFO treatment. **B.** Comparing ROS levels, using the CellROX-green sensor (see Methods), in neurons subjected to cold and the indicated drugs, relative to non-treated cells incubated at 37°C. n= 3 experiments; p values were calculated by ANOVA plus post hoc Tukey test. * indicates p < 0.05; and “ns” not significant. Drugs were administered immediately before transferring cells to cold. Error bars indicate SEM. **C.** Viability of murine neurons, subjected to cold and the indicated antioxidants, examined by staining with propidium iodide. “Edaravone” acts against oxygen and hydroxyl radicals, and inhibits lipid peroxidation/lipoxygenase pathways; “NAC”, N-Acetyl-Cysteine, a glutathione precursor; “TEMPOL” possesses superoxide dismutase (SOD) and catalase (CAT) mimetic properties. Error bars represent SEM. n= 3 experiments; p values between samples were calculated with ANOVA plus Tukey post hoc test was used to compare samples subjected to different drugs. *** indicates p < 0.001. Note that all tested antioxidants enhance cold resistance. **D.** Viability of murine neurons overexpressing Fth1/ferritin heavy chain. Following lentiviral incorporation of EGFP-fused *Fth1*, and a 4 hour incubation in the cold, the survival of transfected neurons was compared to controls; the “mock” neurons were transfected with an "empty" pLJM1-EGFP plasmid. Error bars represent SEM. n= 3 experiments; p values between samples were calculated with ANOVA plus Tukey post hoc test was used to compare samples subjected to different drugs. *** indicates p < 0.001; and ** p < 0.01. Note that neurons overexpressing EGFP-Fth1 survive cold better than control neurons.

If, as expected, decreasing ROS is important for the recovery from cold, treating neurons with antioxidants should provide similar benefit as DFO. To test that, we selected three therapeutic antioxidants: Edaravone ^48–50^, N-acetylcysteine, NAC ^51–53^, and TEMPOL ^54, 55^. Indeed, treating neurons with these drugs strongly enhanced cold survival (Fig. 7C). Finally, we decided to test whether, similar to *ftn-1* overexpression in nematodes, overexpression of its mammalian counterpart, *Fth1*, improves cold survival of neurons. Indeed, we found that *Fth1*-overexpressing neurons survived cold significantly better than mock-transfected neurons (Fig. 7D).

Summarizing, cultured neurons appear to respond to hypothermia in a manner remarkably reminiscent of nematodes. Although Fe(II) was only imaged in neurons, both nematodes and neurons display a transient increase in ROS during rewarming. Moreover, induction of FTN-1/FTH1 enhances cold survival in both models. Presumably, this reflects the capacity of both orthologous proteins for iron detoxification and, consequently, for reducing ROS. Importantly, targeting the free iron-Fe(II)-ROS axis with drugs enhances neuronal cold resistance, suggesting that these and related drugs might prove beneficial in treating hypothermia.

## DISCUSSION

Using a simple, genetically tractable animal model for cold preservation, we uncovered a conserved ability of ferritin to promote cold survival. In this context, FTN-1/FTH1 (ferritin heavy chain) appears to function as an endogenous antioxidant, which counteracts iron-mediated cytotoxicity arising during the recovery from cold. Intriguingly, the elevated expression of *FTH1* has been recognized as a distinctive feature of cold adaptation in hibernating primates, during both daily torpor and seasonal hibernation ^56, 57^. Thus, a mechanism that we turned on artificially in nematodes, through genetic manipulation, may be used actually by some hibernators as a cell-intrinsic mechanism boosting cold resistance. What causes in those animals *FTH1* upregulation remains unknown. However, FoxO3a (related to DAF-16 described here) is upregulated in hibernating squirrels ^58^, arguing for a conserved function of FoxO transcription factors in cold resistance, which might involve the induction of ferritin.

How exactly, and why, cold triggers the accumulation of toxic iron remains to be fully understood. In mammalian epithelial cells, a sizeable fraction of cold-induced free iron was proposed to originate from the microsomal cytochrome P-450 enzymes, which require iron-containing heme as a cofactor ^59^. It was also suggested that the cytosolic free iron causes mitochondrial permeabilization, resulting in apoptosis ^60^. However, as also shown here, treating neurons with BAM15, a mitochondrial uncoupler, suppresses both ROS and cold-induced death ^46^. Assuming that the same treatment reduces iron(II) levels, it may instead suggest the mitochondrial origin of iron toxicity, which agrees with the general view that most cellular ROS is of mitochondrial origin ^61^. Irrespective of its sources, once accumulating, iron(II) can catalyze the formation of ROS, damaging diverse cellular components (like lipids, proteins and nucleic acids), and causing a number of acute and chronic degenerative conditions ^62, 63^.

Crucially, the antioxidant defense is a hallmark of hibernation, being particularly critical during the entry to and exit from hibernation, when oxygen-sensitive tissues, like the brain, are particularly vulnerable to ischemia/reperfusion injury ^4^. Thus, the challenges associated with cooling appear to be, to some extent, similar to those facing the brain in other pathologies. Indeed, the antioxidants employed here have been used to improve outcomes of acute ischemic stroke (Edaravone), hypoxic-ischemic encephalopathy (NAC), and iron-induced cerebral ischemic injury (TEMPOL). Therefore, activating mechanisms protecting cells from cold, or mimicking their effects with drugs, could benefit not only hypothermia patients, but be potentially useful in treating other pathologies, like stroke or neurodegenerative disorders.

## ACKNOWLEDGMENTS

We are grateful to Susan Gasser and the Gasser lab for supporting TP in later stages of her PhD. We thank S. Smallwood and S. Thiry for assistance with mRNA sequencing, L. Gelman and S. Bourke for imaging support, and W. Wendlandt-Stanek for FTN-1 modeling. We thank Collin Ewald, Jacek Kolanowski, Gawain McColl, and Göran Nilsson for discussions and comments on the manuscript. The project POIR.04.04.00-00-203A/16 was carried out within the Team program of the Foundation for Polish Science, co-financed by the European Union under the European Regional Development Fund. RC was also supported by the EMBO Installation Grant No. 3615, the Polish National Science Centre grant 2019/34/A/NZ3/00223, and the Research Council of Norway grant FRIMEDBIO-286499. KS-K and MFi were supported by the National Science Centre grant 2018/31/B/NZ3/03621. Some of the strains were provided by the Caenorhabditis Genetics Center (CGC) funded by the NIH. The Nestin antibody was obtained from the Developmental Studies Hybridoma Bank, created by the NICHD of the NIH, and maintained at the University of Iowa.

## AUTHOR CONTRIBUTIONS

TP performed and analyzed most nematode experiments in Figs. 1–3. JL and KŚ-K performed and analyzed experiments in Figs. 6–7, under the guidance of MFi, who also provided the neuronal stem cell-sphere model. AK and DS performed and analyzed nematode experiments shown in Figs. 4–5; MFr oversaw the ICP-MS experiments. YG analyzed the genomic data, and performed some nematode experiments. RC conceived and supervised the project. RC with other authors wrote the manuscript.

## METHODS

### *C. elegans* handling and genetic manipulation

Animals were grown at 20°C on standard NGM plates, fed with the OP50 *E. coli* bacteria ^65^. All strains used in this study are listed in Table S1. The CRISPR/Cas9 genome editing was used by SunyBiotech to generate the *ftn-1* ferroxidase-dead mutant (allele *syb2550*), and to tag *daf-16* and *pqm-1* (alleles *syb707* and *syb432,* respectively). The latter was achieved through the C-terminal, in-frame insertion of GFP-FLAG (*daf-16*) or mCHERRY-MYC (*pqm-1*). The FTN-1 overexpressing strains (alleles *sybSi67* and *sybSi72*) were generated (by SunyBiotech) using the MosSCI method, utilizing the insertion locus ttTi5605. The *ftn-1* OE constructs were generated using the MultiSite Gateway Technology, and contain circa 2 kb of *dpy-30*, or 1.4 kb of *vit-5* promoter, the genomic *ftn-1* DNA, and 0.7 kb of the *unc-54* 3’UTR. The *sod-5::GFP* strain, GA411, was kindly provided by David Gems.

For RNAi experiments, 1 mM IPTG was added to an overnight culture of RNAi bacteria. 300 μl of bacterial suspension was plated onto agar plates containing 100 μl/ml Carbenicillin and 1 mM IPTG. The L4440 (empty) vector was used as a negative RNAi control. Animals were typically placed on RNAi plates as L1 larvae, and then were grown to day 1 adulthood at 20°C, at which time point they were cold-adapted and scored as described. The RNAi clones used in this study came from either Ahringer or Vidal libraries.

### The assay for *C. elegans* cold survival

Unless stated otherwise, all cold survival experiments were performed in the following way: prior to cold adaptation, animals were grown at 20°C for two generations on OP50. They were then synchronized by bleaching, and L1 larvae were grown until day 1 of adulthood at 20°C. At day 1 of adulthood, they were cold-adapted at 10°C for 2 h, and then shifted to 4°C. Animals were sampled at indicated intervals, and their survival was scored after 24 h recovery at 20°C. Pairwise Wilcoxon signed rank test, in R, was used for statistical comparisons of survival curves between strains. Original counting data and statistical results are included in Table S2.

To examine the sensitivity to ferric ammonium citrate (FAC), animals were grown at 20°C, from L1s to day 1 of adulthood, on different concentrations of FAC, which was added to agar in plates. Animals were then cold-treated and scored for survival as above.

### Poly-A mRNA sequencing

1000 day one adult animals were collected 24 h after cold adaptation. All steps up to Trizol collection were performed at 4°C. Animals were washed 2 times in M9 buffer and snap-frozen in Trizol. Samples were then lysed by freeze/thaw cycles, and RNA extraction proceeded as described before ^66^. Genomic DNA was removed using RNeasy Plus Mini Kit (Qiagen). Quality of RNA was monitored by Bioanalyzer RNA Nano chip (Agilent Technologies). The library was prepared using the TruSeq Library Preparation Kit (Ilumina). Poly-A mRNA was sequenced using a Hiseq 50-cycle single-end reads protocol on a HiSeq 2500 device (Illumina). Raw RNA sequence data were deposited at GEO with accession No. GSE131870.

### Genomic data analysis

FASTQC ^67^ was used to check the quality of the raw sequence data. The reads were mapped to the *C. elegans* genome (Ensembl WBcel235) using STAR ^68^, with default parameters except: outFilterMismatchNmax 3,outFilterMultimapNmax 1, alignIntronMax 15000, outFilterScoreMinOverLread 0.33, outFilterMatchNminOverLread 0.33. Count matrices were generated for the number of reads overlapping with the exons of protein-coding genes using summarizeOverlaps from GenomicFeatures ^69^. Gene expression levels (exonic) from RNA-seq data were quantified as described previously ^70^. After normalization for library size, log2 expression levels were calculated after adding a pseudocount of 8 (y = log2[x + 8]). Genes with 2-fold changes in both replicates were considered significantly differentially expressed. The ChIP bigWig files for PQM-1 and DAF-16 was obtained from ENCODE project ^30^. EnrichedHeatmap ^71^ was used to generate the integrative heatmap.

### RT-qPCR

Around 1000, 1 day-old adult *C. elegans* were collected at 20°C prior to cold adaptation, or at 1 day/ 3 days at 4°C after adaptation, washed 2 times in M9 buffer at the respective temperature, and flash-frozen in Trizol. RNA was isolated as above. 300 ng, or 1000 ng of RNA was used to prepare cDNA with the QuantiTect Reverse Transcription kit (Quiagen), or High-Capacity cDNA Reverse Transcription Kit (Applied Biosystems). cDNA was diluted 1:10 or 1:5 and 5 μl or 2 μl was used with the Light Cycler Syber Green master mix (Roche), or AMPLIFY ME SG Universal Mix (Blirt), and Ct values were calculated using Light Cycler 480 (Roche). *act-1* (actin) was used as the reference gene. Statistical analysis on all of the experiments was performed using the GraphPad/ Prism 8. Statistical method used to calculate P value is indicated in the figure legend. The following primers were used: *act-1* FW: CTATGTTCCAGCCATCCTTCTTGG, *act-1* RV: TGATCTTGATCTTCATGGTTGATGG; *ets-4* FW: CTGAGAACCCGAATCATCCA, *ets-4* RV: TCATTCATGTCTTGACTGCTCC; *ftn-1* FW: CGGCCGTCAATAAACAGATTAACG, *ftn-1* RV: CACGCTCCTCATCCGATTGC; *daf-16* FW: AAAGAGCTCGTGGTGGGTTA, *daf-16* RV: TTCGAGTTGAGCTTTGTAGTCG; *pqm-1* FW: GTGCATCCACAGTAAACCTAATG, *pqm-1* RV: ATTGCAGGGTTCAGATGGAG; *ftn-2* FW: GAGCAGGTCAAATCTATCAACG, *ftn-2* RV: TCGAAGACGTACTCTCCAACTC; *sod-5* FW: ATTGCCAATGCCGTTCTTCC, *sod-5* RV: AGCCAAACAGTTCCGAAGAC.

### Fluorescent imaging of *C. elegans* intestinal nuclei

1 day-old *C. elegans* were anesthetized in 20 μM levamisol and placed on 2 % agar pads. DAF-16::GFP:::FLAG and PQM-1::mCHERRY::MYC were imaged on a spinning disc confocal microscope: Zeiss AxioImager equipped with a Yokogawa CSU-W1 scan-head, 2 PCO Edge cameras, a Plan-Apochromat 40x/1.3 oil objective and two 488 nm and 561 nm laser lines. Laser intensities and exposure times were kept constant for all samples, camera binning was set to 2. Mean fluorescence intensity in intestinal cell nuclei (three per nematode) was quantified manually with FIJI/ImageJ ^72^. The mean fluorescence intensities of each nucleus were averaged and represent one data point for each animal. 10-15 animals were scored per genotype and biological replicate, in total around 40 animals per condition. Statistical analysis was performed using the GraphPad/ Prism 8. Two-tailed, unpaired, t-test was performed to calculate the p value between conditions.

### Fluorescent imaging of SOD-5::GFP

1 day-old adults were anesthetized in 10 mM levamisol and placed on 2 % agar pads. The GFP fluorescence was imaged on Axio Imager.Z2 (Carl Zeiss), equipped with Axiocam 506 mono digital camera (Carl Zeiss), and a Plan-Apochromat 63x/1.40 Oil DIC M27 objective. Images, acquired with the same camera settings, were processed with ZEN 2.5 (blue edition) microscope software in an identical manner, and imported into Adobe Illustrator. 10-15 animals were imaged per time point and biological replicate.

### Oil red O staining and analysis

Oil red O staining was performed as published ^73^. In brief, 0.5 g of Oil Red O powder was mixed in 100 ml isopropanol for 24 h, protected from direct light. This solution was diluted in water to 60 %, stirred O/N, and sterile-filtered using a 0.22 μm pore filter. Between 200-300 day one-old animals were collected with 1 ml of M9 buffer and were washed once with M9. They were fixed in 75 % isopropanol for 15 min with gentle inversions every 3-4 minutes. 1 ml of filtered 60 % ORO was added to the animals after the removal of isopropanol. Staining was performed for 6 h on a shaker with maximum speed, covered with aluminum foil. Stained animals were placed on 2 % agar pads and imaged. Imaging and image analysis were performed as described before ^17^. Briefly, animals were imaged using a wide-field microscope Z1 (Carl Zeiss) using a 10x objective and a color camera AxioCam MRc (Carl Zeiss). RGB images were first corrected for shading in Zen Blue software (Carl Zeiss). Afterwards, images were analyzed using Fiji/ImageJ software suite ^72^, stitched with the Grid/Collection stitching plug-in ^72^, and corrected for white balance. After conversion from RGB to HSB color space, red pixels were selected by color thresholding. A binary mask was created with the Saturation channel and applied to the thresholded image. After conversion to 32-bit, zero pixel values were replaced by NaN. The mean intensity of all remaining pixels was used as a representation of the amount of red staining in the animals (Fiji/ImageJ macro available upon request). 10-15 animals were imaged per genotype and biological replicate. 2 tailed t-test was used to assess significance with Graph Pad/Prism 8.

### *C. elegans* extract preparation

Animals were collected in M9 buffer in a cold room, washed and resuspended in TBS pH 8.0 (2000 worms in 50 μl total volume) with proteinase inhibitors (EDTA-free, Roche) in protein LoBind tubes (Eppendorf). Probes were then homogenized at 4°C in Bioruptor Pico sonicator (Diagenode), using 30 sonication cycles (30 s on/off). After lysis confirmation, by microscopic inspection, probes were centrifuged for 2 h at 21130 *g* at 4°C, and supernatants were transferred to fresh LoBind tubes. Total protein concentration in the soluble fraction was determined by UV absorbance (NanoDrop, Thermo Fisher Scientific). For ICP-MS analysis, *C. elegans* extracts were diluted to 10 μg/μl concentration and transferred to 2 ml glass vials (ALWSCI technologies) with 50 μl glass inserts with bottom spring (Supelco) and kept at 4°C before the analysis.

### Size exclusion chromatography inductively coupled plasma mass spectrometry

All experiments were performed using a ICP-MS‐2030 Inductively Coupled Plasma Mass Spectrometer (Shimadzu, Japan), directly coupled to a Prominence LC 20Ai inert system (Shimadzu, Japan) ^74^. Time-Resolved Measurement (TRM) software for LC–ICP-MS was used for controlling both ICP and LC analytical systems. The ICP-MS operates at 1000 W, with an 8.0 ml min^−1^ argon plasma gas flow, a 0.7 ml min^−1^ Ar carrier gas flow, and a 1.0 ml min^−1^ Ar auxiliary gas flow. The sampling depth was 5.0 mm, and the chamber temp. was set to 3°C. Optimized conditions of the collision cell were −90 V of cell gas voltage, 6.5 V of energy filter voltage, and a 9.0 ml min^−1^ cell gas (He) flow rate. The separation was performed using BioSEC-5, 300A, 5 μm, 4.6×300 mm (Agilent, USA) column using 200 mM ammonium nitrate (99.999% trace metal basis, Sigma Aldrich) pH 8.0 (adjusted by NH_4_OH, Sigma Aldrich, Merck group, Poland) as the mobile phase with a flow rate 0.4 ml min^−1^ and run time 10 min. In all measurements, a 10 μl sample loop was used. For iron content quantification in fractions after separation, ferritin (iron) from equine spleen – Type I standard (Sigma Aldrich) was diluted to 100, 250, 500, 1000 and 2000 μg L^−1^ total metal concentration in mobile phase solution and used to create a standard calibration curve. Iron content in each fraction was normalized to peak area on the chromatogram. Total iron concentration was determined by direct sample injection (LC-ICP-MS) and quantification based on iron standard solution (Sigma Aldrich). The standard calibration curve was created using the same iron concentrations as for SEC-ICP-MS method.

### Suspension culture of mouse neuronal stem cells (NSC) using neuronal cell spheres

Entire heads of fetal mouse (C57BL/6; gestation day between E9-11) were isolated and the tissue was fragmented into pieces followed by incubation in Trypsin-EDTA (0.05 %) (Thermo Fisher Scientific, Waltham, USA) for 15 min at 37°C. The tissue was subsequently transferred to DMEM/10 % FCS and triturated by pipetting up and down into single-cell suspension. The cell suspension was transferred on adherent, uncoated tissue culture plates. After the 3 hour incubation in 5 % CO_2_ at 37°C, the residual differentiated and non-neuronal cells readily attached to the bottom of the plate and the floating neuronal stem cells were collected. The neuronal cells were transferred on to low-adhesive 6-well plates, coated with Poly-HEMA (Poly 2-hydroxyethyl methacrylate; Sigma-Aldrich, St. Louis, USA) using DMEM medium (Thermo Fisher Scientific), supplemented with F-12 (Thermo Fisher Scientific), B-27 (Thermo Fisher Scientific), 100 ng/ml basic fibroblast growth factor (FGF-2, ORF Genetis, Kopavogur, Iceland), 100 ng/ml epidermal growth factor (EGF, ORF Genetis) and 5 μg/ml heparin (Sigma-Aldrich). After 1 day in culture (5 % CO_2_/5 % O_2_), the neuronal cells formed neuronal spheres, which were further cultured and passaged weekly using tissue chopper ^75^.

### Differentiation of NSC to noradrenergic neurons

Neuronal spheres were differentiated towards noradrenergic neurons using available protocols ^76^. The spheres were dissociated by chopping into small cell aggregates and plated onto glass coverslips coated with 0.05 mg/ml poly-D-lysine (Thermo Fisher Scientific) and 3.3 μg/ml laminin (Sigma-Aldrich). Cells were incubated for 5 days in Neurobasal Medium, supplemented with B-27 serum-free supplement, penicillin/streptomycin (all Thermo Fisher Scientific), and neurotrophic factors: 50 ng/ml BDNF, 30 ng/ml GDNF (Peprotech, UK), according to a modified protocol described elsewhere ^76^.

### Detection of NSC and mature neuronal markers

Neuronal stem cell and noradrenergic neuronal cell identities were confirmed by PCR-based detection of neuronal stem cell (NSC) gene markers: *Sox2*, *Gbx2*, *Cd-81*, *Cdh1*, *S100b*, *Dach1*, *Pax6*, *Olig1,* or neural differentiation markers: *Cspg4, DβH, Darpp32, Nestin (NES)*. Moreover the neuronal spheres were immunostained for neuronal stem cell markers: Nestin (1: 500; DSHB, Iowa, USA; ^77^, Foxg-1 (1:100; Abcam, Cambridge, UK), Emx1 (1: 100; Millipore, Burlington, USA) and Emx2 (1:100; Abgent, San Diego, USA) and differentiated neurons for Th (1:100; Abcam), S100b (1:100; Abcam), DβH (1:500; Abcam), Darpp32 (1:50; Abcam).

### Subcloning, lentivirus generation and transduction for Fth1 overexpression in mouse noradrenergic-like neurons

The mouse *Fth1* was introduced into neurons using lentiviral pLJM1 vector for EGFP fusion. pLJM1-EGFP was a gift from David Sabatini (Addgene plasmid # 19319; http://n2t.net/addgene:19319; RRID:Addgene_19319) ^78^. It co-expresses EGFP and a puromycin resistance, and allows for the visualization and selection of transductants carrying N-terminally EGFP-tagged *Fth1*. After RNA isolation from mouse brain tissue, the cDNA template was synthesized using NEBNext Second Strand Synthesis Enzyme Mix for ds cDNA and Phusion® High-Fidelity DNA Polymerase (M0530, NEB). For constitutive expression, the coding sequence of *Fth1* was PCR-amplified (using ProtoScript II Reaction/Enzyme Mix by New England BioLabs, Ipswich, USA).

The following primers with specific Gibson’s overhangs were used: mFth1overexpGibson_F (TCCGGACTCAGATCTCGAGCTCAAGCTTCGATGACCACCGCGTCTCCCTCG), mFth1overexpGibson_R (GATGAATACTGCCATTTGTCTCGAGGTCGAGTTAGCTCTCATCACCGTGTCCC), and cloned into EcoRI-digested lentiviral pLJM1::EGFP vector. Cloning and DNA preparations were done using NEB® Stable Competent *E. coli* (C3040H), according to Gibson Assembly® protocol (NEB).

Lentiviral particles were assembled using a third-generation packaging system. The plasmid pLJM1::EGFP::Fth1 or “empty” pLJM1::EGFP vector, pMDL, pMD2.G and pRSV/REV were mixed (3:2:1:0.8), and human embryonic kidney 293 cells (HEK 293T) (3 × 10^6^ cells seeded on T75 flask one day before) were transfected using a calcium phosphate protocol. Pseudoviral particles in neuronal maintenance medium were collected at 48 and 72 h post-transfection and filtered through a 45 μm filter. The aliquots were snap-frozen and stored at −80°C. Transduction in neurons was done by replacing culture medium with one enriched in lentiviral particles (pLJM1::EGFP::Fth1 or empty pLJM1::EGFP), collected before and supplemented with polybrene (5 μg/mL). After 24 h incubation, medium was discarded and a new one with lentiviral vector was added for subsequent 24 h. At 48 h after transduction, the medium was replaced with fresh viral-free medium and, at 96 h post-transduction, selection with puromycin was initiated for further 48 hours.

### Cold treatment of noradrenergic-like neurons

The assay was established using two independent humidified airtight cell culture incubators. One water-jacketed type incubator was additionally equipped with cooler unit (10°C) and the other incubator was set to 37°C. Both contained atmosphere control, which was set to 5 % CO_2_/5 % O_2_. If not stated otherwise, differentiated neuronal cultures were placed in a 10°C-incubator for 4 hours and then returned to the 37°C-incubator for additional 24 hours of rewarming. Such cooling/rewarming paradigm demonstrated a statistically relevant rise of cell death as early as 4 h into cooling, which was used in subsequent assays. To evaluate neuroprotective effects of compounds, neuronal culture medium was replaced with a Neurobasal medium, without neurotrophic factors supplemented with 100 μM deferoxamine (DFO concentration determined based on dose curve at 10°C) (Sigma-Aldrich), 100 nM BAM15 (Tocris, Bristol, UK), 1:500 dilution of protease inhibitor cocktail III PI (Sigma-Aldrich). All compounds were provided as a single or as a combined treatment. The effects of antioxidants were tested by supplementation of neuronal maintenance medium with 50 μM Edaravone (Sigma-Aldrich), 50 μM TEMPOL (Sigma-Aldrich), or 10 μM N-Acetyl-L-cysteine NAC (Sigma-Aldrich), following procedures described above. Drugs concentration was determined based on dose curves at 10°C.

### Propidium iodide staining

After 4 hours of cooling at 10°C, and additional 24 hours of rewarming at 37°C, neurons cultured on glass coverslips were incubated in the presence of 10 μg/ml propidium iodide (PI) (Cayman Chemical, Ann Arbor, USA) diluted in phosphate-buffered saline (PBS), and costained with 1 μg/ml Hoechst 33342 (Life Technologies) for 25 min at 37°C. Cells were then fixed in ice-cold 4 % buffered formaldehyde for 15 min, washed twice in PBS and placed in histology mounting medium (Sigma-Aldrich) on a glass slide. The prepared material was imaged using fluorescence microscope (Leica DMI 4000B, Germany) and LAS X SP8 software. Counting of total cells (blue nuclei) and necrotic cells (red-PI positive and round) was performed on 2-3 images from 3 coverslips as replicates. Collected data were statistically analyzed using Prism software (version 6.01 for Windows, La Jolla, CA).

### CellROX-green staining of ROS production

To demonstrate whether cold-stabilizing drugs inhibit intracellular ROS levels, neurosphere-derived neurons were incubated with cell-permeable dye, CellROX Green Reagent (Life Technologies, Carlsbad, USA), at the final concentration of 5 μM, according to the procedure described elsewhere ^46^. Green fluorescence was emitted after dye binding to DNA, only upon its oxidation. In brief, murine neuronal cells were differentiated in 24-well plates on glass coverslips. Control cells (group 1) were maintained only at 37°C (non-cold control).

Other cells (group 2) were exposed for 4 hrs to 10°C, in the presence or absence of the following drugs: 100 μM DFO, 100 nM BAM15 and 1:500 dilution of protease inhibitor cocktail (PI). Subsequently, after 5 min rewarming at room temperature on the bench, ROS accumulation in neurons was assessed for both groups, by staining with fluorogenic Cell-ROX green reagent in the dark for 30 min at 37°C. Additionally, before cooling, reference neurons (for time 0) were labeled to detect fluorescent signal in initial precooling cultures. Z-stack well-focused confocal images at 0.55 μm intervals in the z-axis of 4 culture areas per each treatment and condition were collected for both groups. Maximal intensity projections of the Z stack images were produced for data analysis. Microscopy images were taken using a Leica TCS SP5 confocal microscope and LAS X SP8 software. The change in ROS production at 10°C incubated cells vs cells maintained only in 37°C was calculated using the following formula: (F_2_ – F_1_)/F_0_ x 100%. The CellROX-green fluorescence of non-cold control cells (F_1_; mean intensity) was subtracted from CellROX-fluorescence at the end of cold treatment (F_2_; mean intensity). While F_0_ stands for the initial mean CellROX-green fluorescent intensity of culture areas before 10°C cooling.

### Determination of iron(II) with FeRhoNox-1

Intracellular iron levels were measured according to the manufacturer’s protocol. Cells were cultured in a glass-bottom dish, and exposed to indicated agents in the cold. Next, cells were rinsed twice with HBSS, then 5 μM FeRhoNox™-1 solution (Goryo Chemical, Inc., Sapporo, Japan) was added and incubated in the dark at 37 °C for 1 h, and then washed twice with HBSS. To track changes in Fe^2+^ over time, after cooling neurons at 10 degrees, the probe signal was recorded at 1, 4 and 8 hours of rewarming. In turn, to examine iron(II) right after cooling (0 hour), incubation with the reagent was completed at the end of the 4-hour cold exposure. The FeRhoNox signal was visualized using a confocal microscope (Leica TCS SP5, Germany) and LAS X SP8 software. FeRhoNox-1 was excited at 543 nm and measured at 570 nm. Fluorescence intensity of Z-stacked confocal images of neuronal culture (maximal intensity projection of 7 image z-stacks at 0.55 μm intervals in the z-axis) was analyzed using ImageJ.

### Neurite tracing

Cultured neurons with, or without, DFO or BAM15/PI/DFO were fixed with 4 % paraformaldehyde, permeabilized and washed with 0.1 % Triton X-100 in phosphate-buffered saline, and stained by antibody against NEFH. NEFH^+^ neurite paths were traced with the ‘Simple Neurite Tracer’ plugin ^79^, ImageJ, using Z-stacked confocal images of neuronal culture (maximal intensity projection of 7 image z-stacks at 0.55 μm intervals in the z-axis). Cumulative frequency plots of neurite lengths, for each experimental group, were built using GraphPad Prism version 6.01 for Windows, GraphPad Software, La Jolla California USA.

