## Supplemental Information for "Surviving hypothermia by ferritin-mediated iron detoxification"

Pekec et al., Figure S1

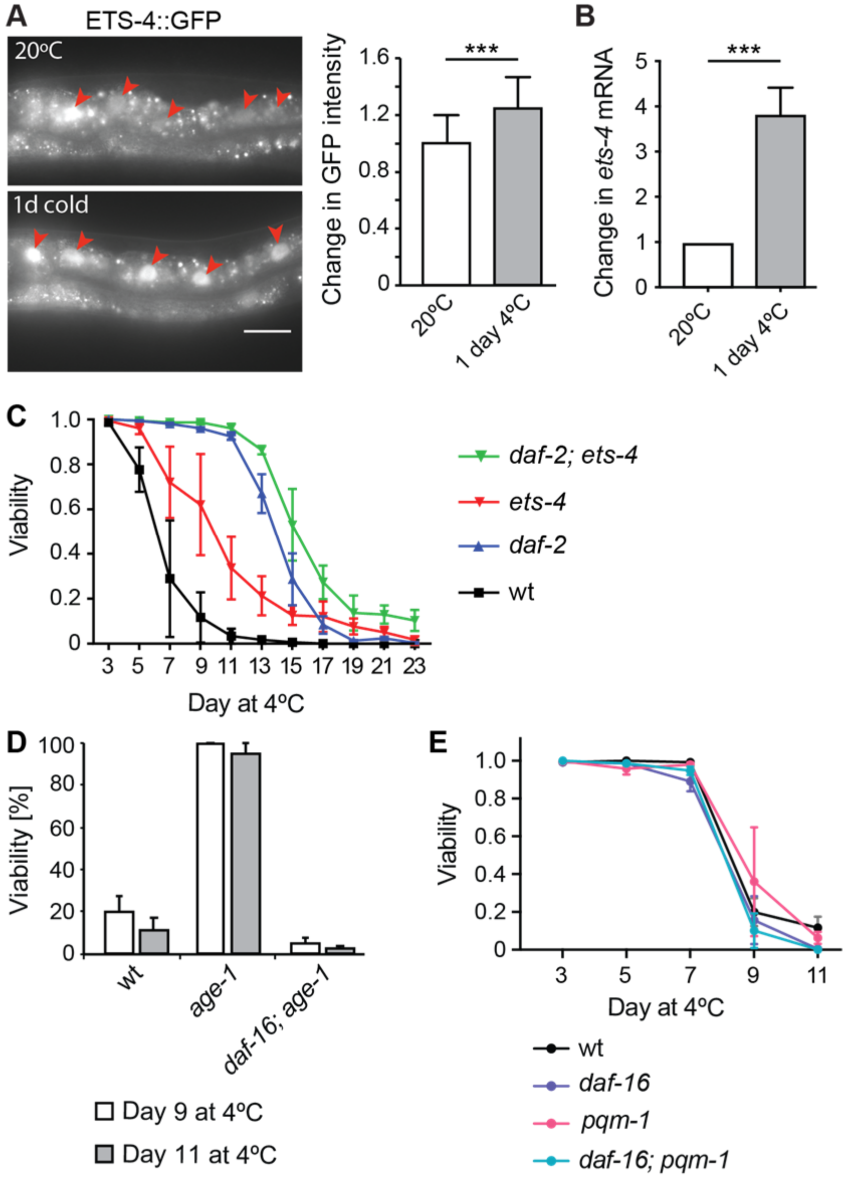

**Figure S1. Cold-mediated upregulation of ETS-4, and the relationship between *ets-4(-)* and insulin pathway mutants.**

**A.** ETS-4::GFP *(rrr45)* expression in the cold. 1 day-old adults were treated according to Fig. 1A. Representative images, on the left, show enhanced GFP fluorescence in the intestinal nuclei (arrowheads). Scale bar: 20 µm. The corresponding quantification is on the right. Student t test was used in statistical analysis. Error bars show standard deviation (SD). ***; $p < 0.001$.

**B.** The levels of *ets-4* mRNA, normalized to *act-1* mRNA, were measured (by RT-qPCR) in 1 day-old adult wild-type animals at 20°C, and after one day at 4°C, without rewarming. P value was calculated using an unpaired student t-test (n= 3). Error bars represent SEM. ***; $p < 0.001$.

**C.** Survival of animals, of the indicated genotypes, subjected to cold as in 1A. Note that *ets-4(rrr16)*, *daf-2(e1370)* and *daf-2(e1370); ets-4(rrr16)* double mutants survived cold far better than wt. Combining *daf-2(e1370)* and *ets-4(rrr16)* mutations only slightly improved cold resistance of the *daf-2(e1370)* single mutant. Error bars represent SEM. n= 3; 324-668 animals were scored per time point.

**D.** Survival of animals, of the indicated genotypes, subjected to cold. Wt, *age-1(hx546)*, and *daf-16(mu86); age-1(hx546)* animals were treated as in 1A. The *age-1(-)* mutant recovered from cold much better than wt, while there was no obvious difference between the double *daf-16(-); age-1(-)* mutant and wt (n= 3; 200-300 animals were scored per time point). Error bars represent SEM.

**E.** Survival of animals, of the indicated genotypes, subjected to cold. Wt, *daf-16(mu86)*, *pqm-1(ok485)*, and the double *daf-16(mu86); pqm-1(ok485)* mutants were cold-adapted (n= 3; 200-300 animals were scored per time point). Error bars represent SEM. There was no obvious difference between any of the examined animals.

Pekec et al., Figure S2

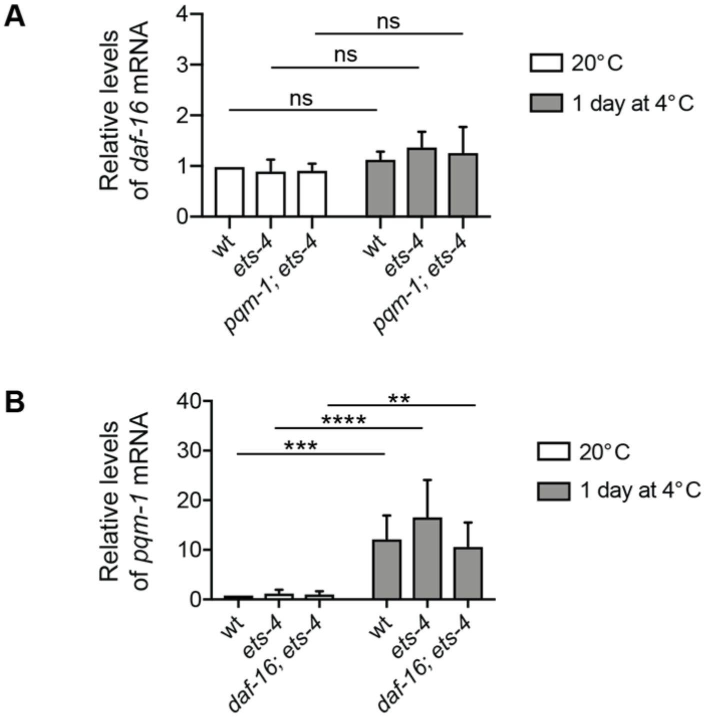


**Figure S2. Evaluation of *daf-16* and *pqm-1* mRNAs levels in the cold.**

**A.** Animals were collected at 20°C, before adaptation, and after 1 day at 4°C, as in 1A. The levels of *daf-16* mRNA, measured by RT-qPCR, were normalized to *act-1* mRNA, and are shown relative to the *daf-16* mRNA level in wt at 20ºC. Wt and *ets-4(rrr16)* mutants were collected at 20°C, and after one day at 4°C (n= 5). P values were calculated using 2-way ANOVA. Error bars represent SEM. “ns” = not significant.

**B.** Relative *pqm-1* mRNA levels were measured as in A. Wt, *ets-4(rrr16)*, and *daf-16(mu86); ets-4(rrr16)* animals were collected at 20°C, and after one day at 4°C (n= 5). P values were calculated using 2-way ANOVA. Error bars represent SEM. ** $p > 0.01$; *** $p > 0.001$; **** $p > 0.0001$.

Pekec et al. Figure S3

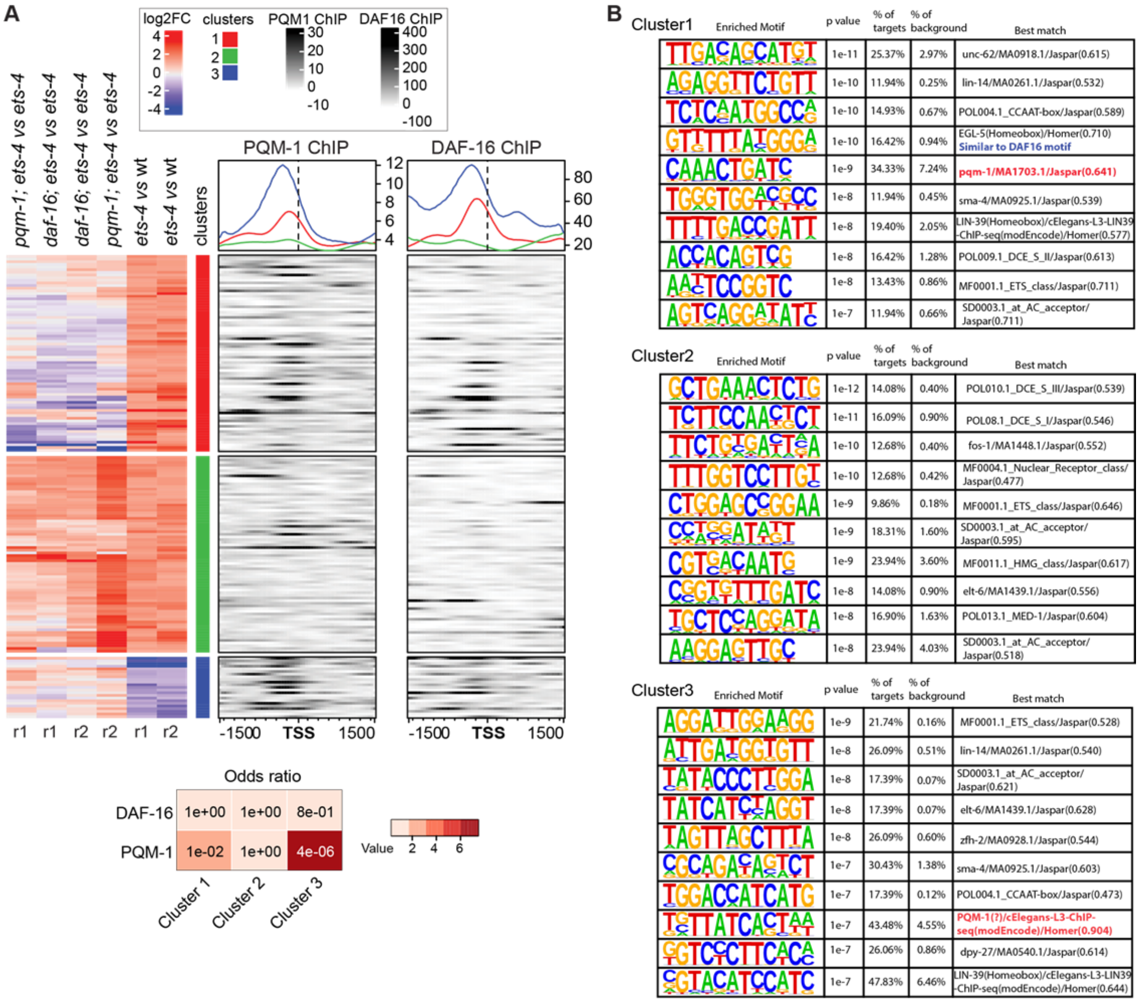


**Figure S3. Identification of genes potentially promoting cold survival.**

**A.** Transcriptome analysis, by RNA-seq, performed on animals of the indicated genotypes, as described in detail in the Methods. Strains used: wt, *ets-4(rrr16)*, *daf-16(mu86); ets-4(rrr16),* and *pqm-1(ok485); ets-4(rrr16)*. The animals, treated according to 1A, were collected at day 1 at 4ºC. Left: Integrative heat map showing $\log_2$ fold changes in gene expression between the indicated strains. “r1 and r2” indicate biological replicates. Each line represents one gene. Automated clustering showed three distinct clusters: cluster 1 (red) includes genes up-regulated in the *ets-4* animals (compared to wild type), and not changing or downregulated in the double mutants (compared to *ets-4*). Cluster 2 (green) includes genes upregulated across all samples. Cluster 3 (blue) includes genes that are downregulated in the *ets-4(-)* animals, and not changing or up-regulated in the double mutants. Right: Using the ChIP ENCODE data [1], we examined the binding of DAF-16 and PQM-1 around the transcription start sites (TSSs) of genes shown on the left. The line graphs above (colored according to the clusters) illustrate enrichments for DAF-16 or PQM-1 binding, within each cluster, around the TSS. Note that both DAF-16 and PQM-1 tend to bind the promoters of cluster 1 (red) and 3 (blue) genes, but not the cluster 2 (green) genes. The heatmap below shows a hypergeometric test of overlaps between 3 clusters of genes from A, and PQM-1 or DAF-16 targeted genes. Color-coded p values are shown.

**B.** De novo motif enrichment analysis of each gene cluster in A. The enrichment analysis were performed with HOMER [2], for each cluster with the following parameters: -start -1500 -end 1500 -p 6. All genes in the genome were used as background. Top 10 enriched motifs per gene cluster are shown, other matched motifs are not shown.

Pekec et al., Figure S4

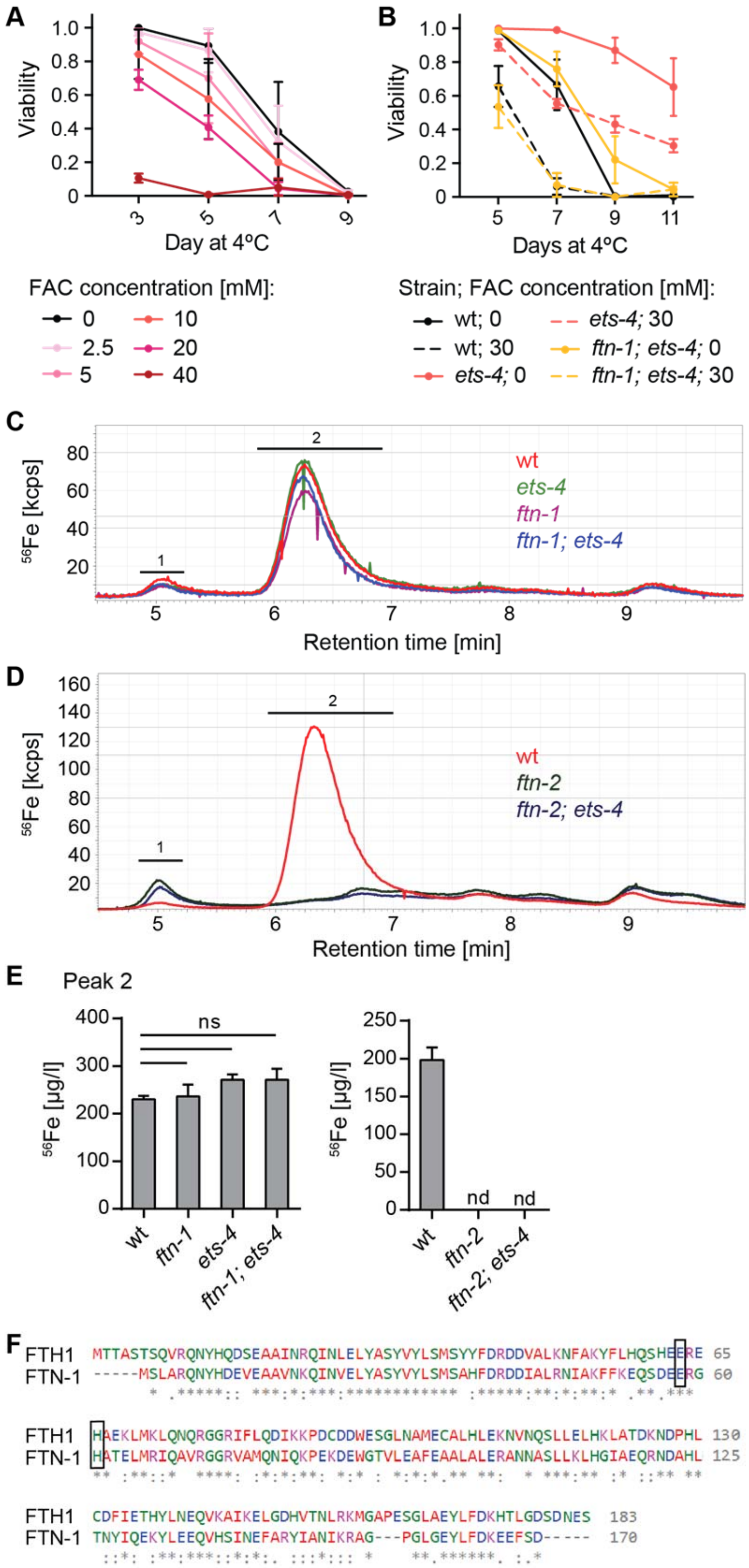

**Figure S4. Characterization of FTN-1 properties.**

**A.** Wt animals were exposed to excess iron (in the form of ferric ammonium citrate, FAC), provided in agarose plates. Animals were grown on FAC-supplemented plates from the L1 larval stage to day one adulthood, prior to cold adaptation as described in 1A. (n= 3; 200-300 animals were scored per condition). Error bars represent SEM.

**B.** Viability of animals, of the indicated genotypes, subjected to cold as in 1A and exposed to excess iron (FAC). Error bars represent SEM. n= 3; 300–500 animals were scored per time point.

**C-D.** Native soluble iron-binding species separated and detected by SEC-ICP-MS. Iron is mostly associated with high molecular weight complexes (peak 1) and ferritin (peak 2). Note that FTN-2, but not FTN-1, significantly contributes to iron binding.

**E.** Integration of the second chromatographic peak corresponding to ferritin, calculated relative to ferritin standard. Only loss of *ftn-2* decreased iron association within peak 2 (ferritin). Strains used: wild type, *ets-4(rrr16)*, *ftn-1(ok3625)*, *ftn-1(ok3625); ets-4(rrr16)*, *ftn-2(ok404)*, *ftn-2(ok404)*; *ets-4(rrr16)*. Unpaired two-tailed t-test was used to calculate the p value, "ns" = not significant, "nd" = not detected. Error bars represent SEM, n= 3.

**F.** Amino acid sequence alignment of *H. sapiens* ferritin heavy chain 1 (FTH1; NCBI accession number NP_002023.2) and *C. elegans* ferritin (FTN-1: NCBI accession number NP_504944.2). The alignment was based on the multiple sequence alignment software Clustal Omega [3]. Residues mutated in *ftn-1*, yielding a ferroxidase-dead protein, are boxed. Asterisks (*) indicate fully conserved residues, colons (:) residues with strongly similar properties, and periods (.) residues with weakly similar properties.

Pekec et al., Figure S5

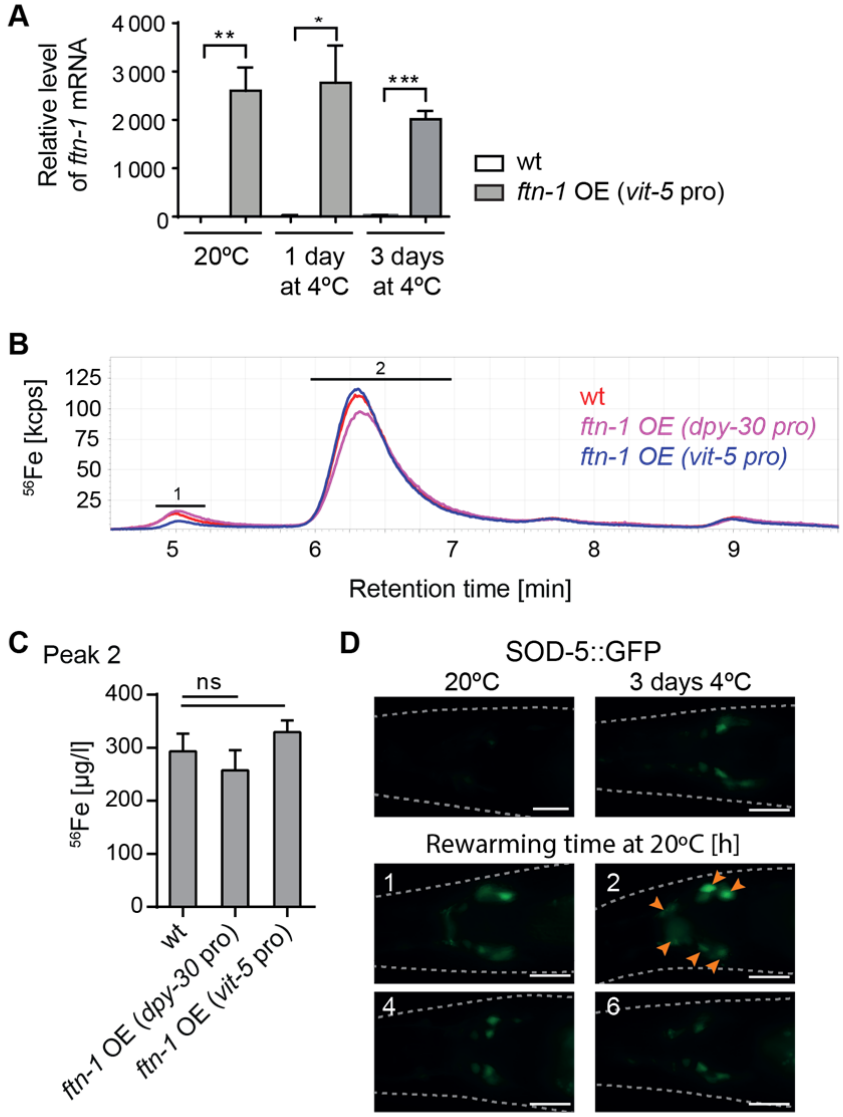


**Figure S5. Overexpressed FTN-1 counteracts the expression of SOD-5, a ROS-induced superoxide dismutase, during rewarming.**

**A.** The levels of *ftn-1* mRNA, measured by RT-qPCR, in animals of the indicated genotypes. 1 day-old adults were collected at 20°C before the cold adaptation, and after one and three days at 4°C without rewarming at 20°C. The mRNA levels were normalized to *act-1* mRNA. At each time points, the values were then normalized to the wt at 20°C. Note greatly elevated levels of *ftn-1* in the strain overexpressing it (OE) from the *vit-5* promoter (*vit-5* pro).

Error bars represent SEM. n= 3; p values were calculated using unpaired Student t-test. * indicates $p < 0.05$; ** $p < 0.01$; *** $p < 0.001$.

**B.** Native soluble iron-binding species separated and detected by SEC-ICP-MS like in S4C-D. Iron is mostly associated with high molecular weight complexes (peak 1) and ferritin (peak 2). Note that FTN-1 overexpression (from either *vit-5* or *dpy-30* promoter) contributes little or nothing to iron sequestration.

**C.** Integration of the second chromatographic peak, corresponding to ferritin, calculated relative to ferritin standard. FTN-1 overexpression did not have any obvious effect on decreased iron association within peak 2 (ferritin). Unpaired two-tailed t-test was used to calculate the p value, “ns” = not significant, “nd” = not detected. Error bars represent SEM, n= 3.

**D.** Representative fluorescence micrographs, showing expression of SOD-5::GFP fusion protein [4] in the head region, in 1 day-old adults subjected to cold. The animals were exposed to cold for 3 days, transferred back to 20°C, and examined during rewarming in a time-course fashion. Images were taken at indicated times and temperatures. Note that a peak of SOD-5::GFP was observed in head neurons around 2 hours of rewarming at 20°C. Animals are outlined with grey, dashed lines. Scale bar: 20 μm.

Pekec et al. Fig. S6

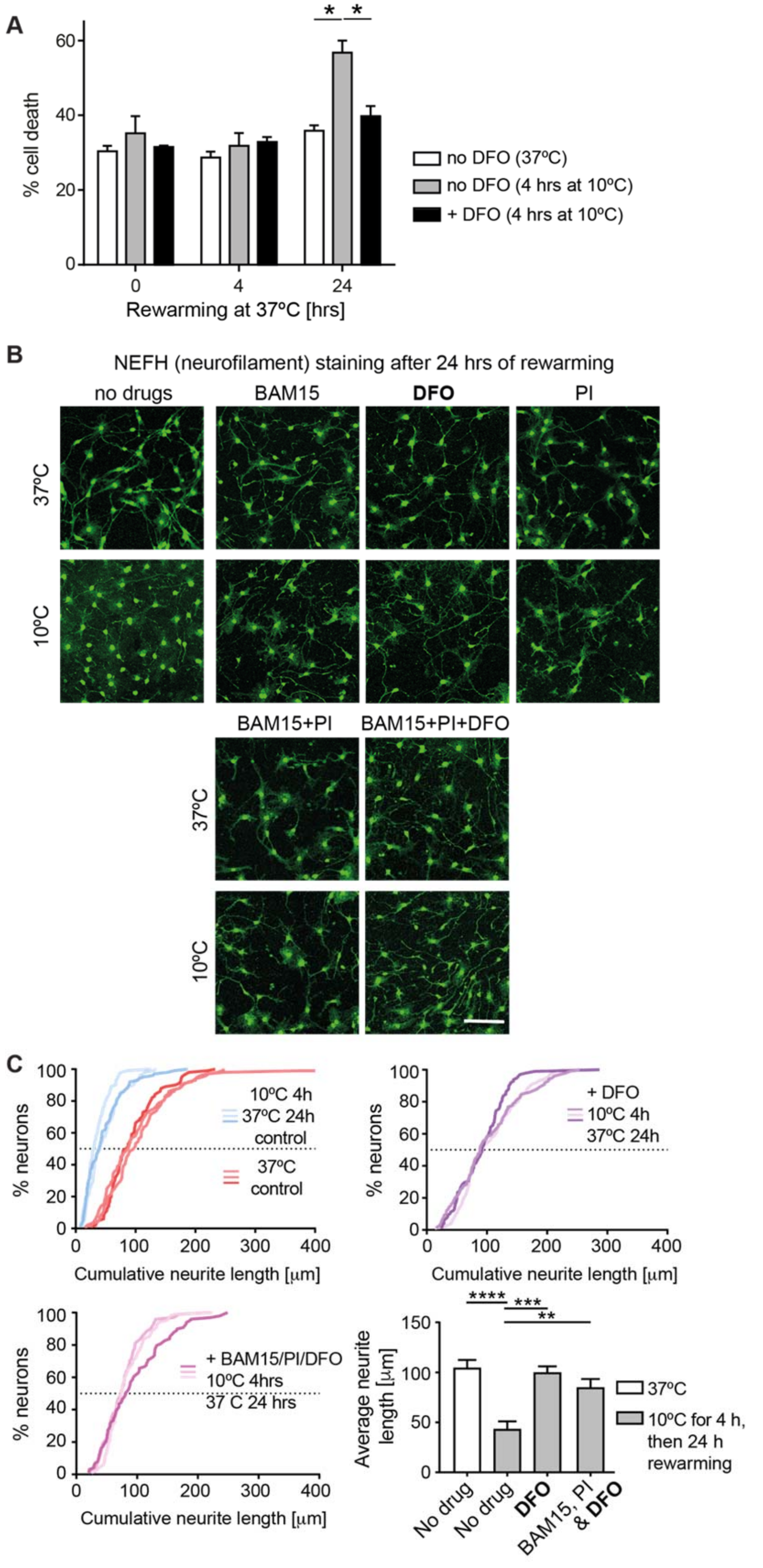


121

**Figure S6. Exposure to cold induces neuronal degeneration during rewarming.**

**A.** Viability of murine neurons, subjected to cold +/- DFO, was examined during rewarming at 37ºC, by staining with propidium iodide (see Methods). Error bars represent SEM. n= 3 experiments; p values were calculated with Student's t-test; * $p < 0.05$. Note that neurons began dying during rewarming, which was prevented by the treatment with DFO.

**B.** Representative confocal images of neurons, subjected to cold and the indicated drugs, and stained, by immunofluorescence, for NEFH to visualize neurites after 24 h rewarming at 37ºC. As in 6B, the neurites, which in control neurons degenerate upon cold exposure, were stabilized by adding DFO, similar to BAM15, PI, or the combinations of drugs. Note that, in contrast to 6B, here the cells were examined after one day of rewarming. The neurites appeared well-preserved, arguing that the protective effects of drugs are long-lasting. Scale bar: 40 μm.

**C.** Quantifications of neurite lengths corresponding to B. The cumulative plots (see Methods) compared neurite lengths in cells treated as indicated. Each curve corresponds to one experimental replicate. The bar graph (bottom right) compares average neurite lengths. Neuroprotective effects of the applied drugs, or DFO alone, were still noticeable following the 24 h rewarming. n= 3 experiments from independent differentiation neuron groups; ANOVA plus post hoc Tukey test for multiple cold-exposed groups, ** $p > 0.01$; *** indicates $p < 0.001$; **** $p > 0.0001$. Error bars indicate SD.

Pekec et al. Fig. S7

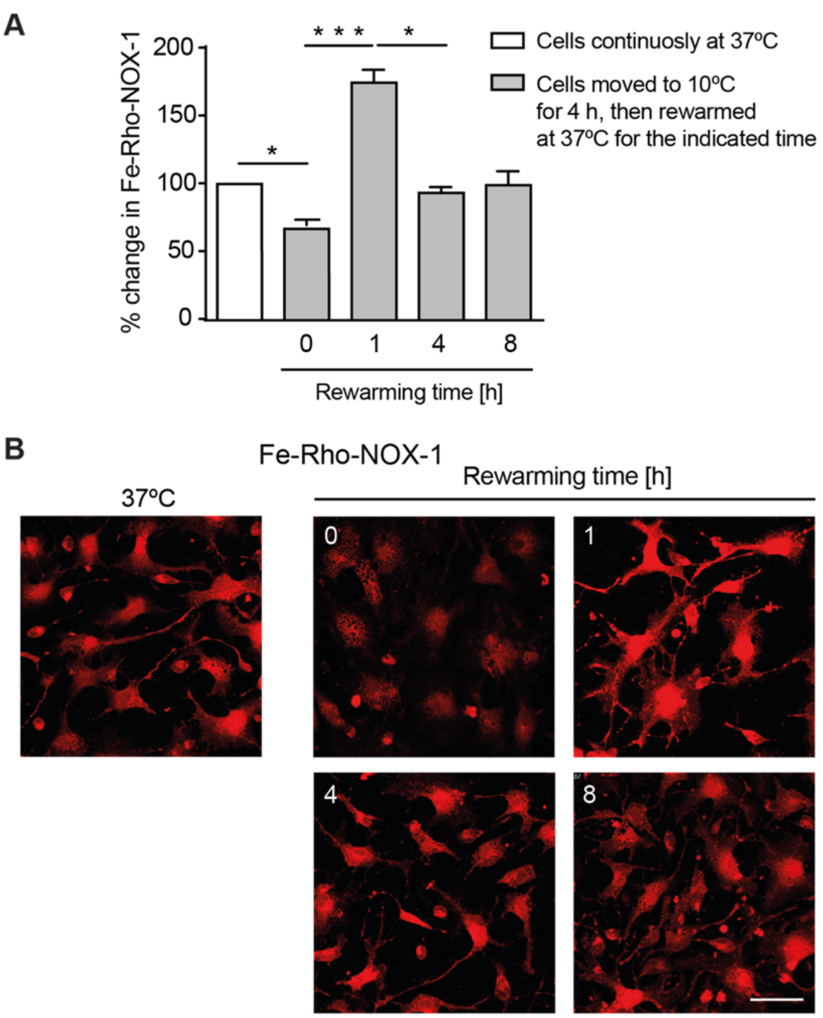


**Figure S7. Lowering iron has a lasting protective effect on neural integrity.**

**A.** Levels of iron(II), observed at different times during rewarming, detected with FeRhoNox-1 in murine neurons. Quantifications of changes in FeRhoNox-1 fluorescence are relative to non-treated cells incubated at 37ºC. n= 3; p values were calculated by ANOVA plus post hoc Tukey test. *** indicates $p < 0.001$; and * $p < 0.05$. Error bars represent SEM. Note the peak of iron(II) shortly into rewarming.

**B.** Representative micrographs of FeRhoNox-1 fluorescence in neurons at the indicated temperatures and times. Rewarming was at 20ºC, following a 4-hour incubation at 10ºC. Scale bar: 50 µm. Note the peak of FeRhoNox-1 fluorescence, corresponding to $Fe^{2+}$, around 1 hour of rewarming.

**Table S1. The *C. elegans* strains used in this work.**

| Genotype | **CGC**/RAF/**Other** |
|---|---|
| *age-1(hx546) II.; ets-4(rrr16) X.* | 2169 |
| *age-1(hx546) II.* | **TJ1052**/1891 |
| *daf-16(mu86) I.; age-1(hx546) II.* | 2150 |
| *daf-16(syb707) I.* | 5010 |
| *daf-16(mu86) I.* | **CF1038**/1660 |
| *ets-4(rrr16) X.* | 1758 |
| *daf-16(mu86) I.; ets-4(rrr16) X.* | 2107 |
| *pqm-1(ok485) II.; ets-4(rrr16) X.* | 2106 |
| *ftn-1(ok3625) V.* | **RB2603**/2162 |
| *pqm-1(ok485) II.* | **RB711**/2104 |
| *daf-16(mu86) I.; pqm-1(ok485) II.* | 2105 |
| *pqm-1(ok485) II.; ets-4(rrr16) X.* | 2033 |
| *pqm-1(syb432) II.* | 2156 |
| *pqm-1(syb432) II.; ets-4(rrr16) X.* | 2157 |
| *rege-1(rrr13) I.* | 5018 |
| *rege-1(rrr13) I.; ets-4(rrr16) X.* | 1759 |
| *daf-16(syb707) I.; ets-4(rrr16) X.* | 5054 |
| *daf-16(mu86) I.; pqm-1(ok485) II.; ets-4(rrr16) X.* | 5062 |
| *ftn-1(ok3625) V.; ets-4(rrr16) X.* | 5063 |
| *daf-2(e1370) III.* | **CB1370**/1173 |

| | |
|---|---|
| *daf-2(e1370) III.; ets-4(rrr16) X.* | 5096 |
| *sybSi67[Pdpy-30::ftn-1::unc-54 3'UTR] II.;unc-119(ed3) III.* | 5069/**PHX1798** |
| *sybSi72[Pvit-5::ftn-1::unc-54 3'UTR] II.;unc-119(ed3) III.* | 5071/**PHX1920** |
| *ftn-2(ok404) I.* | **RB668**/5085 |
| *ftn-2(ok404) I.; ets-4(rrr16) X.* | 5093 |
| *ftn-1(syb2550) V.* | 5118 |
| *ftn-1(syb2550) V.;ets-4(rrr16) X.* | 5112 |
| *wuIs57[pPD95.77 sod-5::GFP, rol-6(su1006)]* | 5106 |

“CGC” indicates strain numbers deposited in the Caenorhabditis Genetics Center; “RAF” strain numbers in the Ciosk lab collection; "Other" strains obtained from other sources.

**Table S2. Analysis of the *C. elegans* cold-survival experiments.**
