## Supplemental Tables for "Surviving hypothermia by ferritin-mediated iron detoxification"

| | 1st replicate | | 2nd replicate | | 3rd replicate | | | | | |
|---|---|---|---|---|---|---|---|---|---|---|
| **wt** | | | | | | | | | | |
| Days at 4°C | Total | Survival | Total | Survival | Total | Survival | Total animals | Average Survival | St.Dev. (+/-) | SEM (+/-) |
| 3 | 68 | 0.99 | 86 | 1.00 | 75 | 0.97 | 229 | 0.9862 | 0.0134 | 0.0077 |
| 5 | 75 | 0.99 | 125 | 1.00 | 63 | 1.00 | 263 | 0.9956 | 0.0077 | 0.0044 |
| 7 | 93 | 0.72 | 117 | 0.12 | 90 | 0.33 | 300 | 0.3911 | 0.3045 | 0.1758 |
| 9 | 117 | 0.06 | 111 | 0.02 | 58 | 0.03 | 286 | 0.0374 | 0.0211 | 0.0122 |
| 11 | 119 | 0.00 | 124 | 0.00 | 84 | 0.00 | 327 | 0.0000 | 0.0000 | 0.0000 |
| ***ets-4(rrr16)*** | | | | | | | | | | |
| Days at 4°C | Total | Survival | Total | Survival | Total | Survival | Total animals | Average Survival | St.Dev. (+/-) | SEM (+/-) |
| 3 | 81 | 1.00 | 125 | 1.00 | 78 | 1.00 | 284 | 1.0000 | 0.0000 | 0.0000 |
| 5 | 91 | 0.98 | 126 | 1.00 | 73 | 1.00 | 290 | 0.9927 | 0.0127 | 0.0073 |
| 7 | 83 | 0.98 | 105 | 0.80 | 67 | 0.82 | 255 | 0.8656 | 0.0961 | 0.0555 |
| 9 | 95 | 0.83 | 104 | 0.45 | 65 | 0.34 | 264 | 0.5407 | 0.2583 | 0.1491 |
| 11 | 105 | 0.58 | 109 | 0.17 | 77 | 0.39 | 291 | 0.3816 | 0.2034 | 0.1175 |
| ***rege-1(rrr13)*** | | | | | | | | | | |
| Days at 4°C | Total | Survival | Total | Survival | Total | Survival | Total animals | Average Survival | St.Dev. (+/-) | SEM (+/-) |
| 3 | 59 | 0.34 | 103 | 0.57 | 70 | 0.17 | 232 | 0.3611 | 0.2016 | 0.1164 |
| 5 | 146 | 0.03 | 136 | 0.06 | 72 | 0.07 | 354 | 0.0542 | 0.0181 | 0.0104 |
| 7 | 129 | 0.02 | 102 | 0.06 | 57 | 0.00 | 288 | 0.0248 | 0.0305 | 0.0176 |
| 9 | 80 | 0.00 | 97 | 0.03 | 69 | 0.00 | 246 | 0.0103 | 0.0179 | 0.0103 |
| 11 | 105 | 0.00 | 126 | 0.00 | 81 | 0.00 | 312 | 0.0000 | 0.0000 | 0.0000 |
| ***rege-1(rrr13); ets-4(rrr16)*** | | | | | | | | | | |
| Days at 4°C | Total | Survival | Total | Survival | Total | Survival | Total animals | Average Survival | St.Dev. (+/-) | SEM (+/-) |
| 3 | 79 | 0.99 | 123 | 1.00 | 70 | 1.00 | 272 | 0.9958 | 0.0073 | 0.0042 |
| 5 | 92 | 0.97 | 93 | 1.00 | 81 | 0.98 | 266 | 0.9809 | 0.0170 | 0.0098 |
| 7 | 90 | 0.81 | 111 | 0.97 | 81 | 0.99 | 282 | 0.9239 | 0.0980 | 0.0566 |
| 9 | 95 | 0.57 | 111 | 0.70 | 53 | 0.55 | 259 | 0.6061 | 0.0843 | 0.0487 |
| 11 | 111 | 0.33 | 95 | 0.54 | 82 | 0.17 | 288 | 0.3470 | 0.1834 | 0.1059 |

Wilcoxon signed rank test

| | *ets-4(rrr16)* | *rege-1(rrr13)* | *rege-1(rrr13); ets-4(rrr16)* |
|---|---|---|---|
| wt | 0.0055 | 0.0055 | 0.0064 |
| *rege-1(rrr13); ets-4(rrr16)* | 1 | 0.0022 | |
| *rege-1(rrr13)* | 0.0022 | | |

| | 1st replicate | | 2nd replicate | | 3rd replicate | | 4th replicate | | | | | |
|---|---|---|---|---|---|---|---|---|---|---|---|---|
| **wt** | | | | | | | | | | | | |
| Days at 4°C | Total | Survival | Total | Survival | Total | Survival | Total | Survival | Total animals | Average Survival | St. Dev. (+/-) | SEM (+/-) |
| 3 | 108 | 0.99 | 105 | 1.00 | 86 | 1.00 | 92 | 1.00 | 391 | 0.9977 | 0.0046 | 0.0023 |
| 5 | 103 | 0.97 | 106 | 1.00 | 150 | 1.00 | 94 | 1.00 | 453 | 0.9927 | 0.0146 | 0.0073 |
| 7 | 116 | 0.97 | 97 | 1.00 | 137 | 0.81 | 120 | 0.70 | 470 | 0.8689 | 0.1396 | 0.0698 |
| 9 | 103 | 0.59 | 124 | 0.11 | 112 | 0.02 | 121 | 0.02 | 460 | 0.1849 | 0.2753 | 0.1376 |
| 11 | 102 | 0.04 | 96 | 0.10 | 149 | 0.00 | 89 | 0.00 | 436 | 0.0358 | 0.0492 | 0.0246 |
| 13 | 105 | 0.00 | 100 | 0.08 | 129 | 0.00 | 154 | 0.00 | 488 | 0.0200 | 0.0400 | 0.0200 |
| 15 | 94 | 0.01 | 146 | 0.00 | 155 | 0.00 | 128 | 0.00 | 523 | 0.0027 | 0.0053 | 0.0027 |
| 17 | 95 | 0.00 | 78 | 0.01 | 135 | 0.00 | 110 | 0.00 | 418 | 0.0032 | 0.0064 | 0.0032 |
| 19 | 96 | 0.00 | 79 | 0.00 | 119 | 0.00 | 129 | 0.00 | 423 | 0.0000 | 0.0000 | 0.0000 |
| 21 | 105 | 0.00 | 90 | 0.00 | 126 | 0.00 | 143 | 0.00 | 464 | 0.0000 | 0.0000 | 0.0000 |
| ***ets-4(rrr16)*** | | | | | | | | | | | | |
| Days at 4°C | Total | Survival | Total | Survival | Total | Survival | Total | Survival | Total animals | Average Survival | St. Dev. (+/-) | SEM (+/-) |
| 3 | 95 | 1.00 | 92 | 1.00 | 97 | 0.99 | 101 | 1.00 | 385 | 0.9974 | 0.0052 | 0.0026 |
| 5 | 80 | 1.00 | 76 | 1.00 | 147 | 1.00 | 79 | 1.00 | 382 | 1.0000 | 0.0000 | 0.0000 |
| 7 | 75 | 0.99 | 77 | 0.99 | 154 | 0.97 | 68 | 1.00 | 374 | 0.9869 | 0.0106 | 0.0053 |
| 9 | 107 | 0.90 | 84 | 0.82 | 141 | 0.63 | 91 | 1.00 | 423 | 0.8375 | 0.1558 | 0.0779 |
| 11 | 80 | 0.71 | 94 | 0.47 | 181 | 0.52 | 66 | 0.91 | 421 | 0.6523 | 0.2010 | 0.1005 |
| 13 | 74 | 0.27 | 81 | 0.52 | 150 | 0.33 | 96 | 0.63 | 401 | 0.4368 | 0.1638 | 0.0819 |
| 15 | 62 | 0.29 | 83 | 0.40 | 135 | 0.24 | 91 | 0.51 | 371 | 0.3576 | 0.1191 | 0.0595 |
| 17 | 80 | 0.35 | 67 | 0.13 | 142 | 0.17 | 67 | 0.42 | 356 | 0.2678 | 0.1377 | 0.0688 |
| 19 | 84 | 0.26 | 75 | 0.37 | 149 | 0.07 | 86 | 0.17 | 394 | 0.2209 | 0.1274 | 0.0637 |
| 21 | 80 | 0.26 | 80 | 0.21 | 133 | 0.15 | 104 | 0.19 | 397 | 0.2044 | 0.0466 | 0.0233 |
| ***age-1(hx546)*** | | | | | | | | | | | | |
| Days at 4°C | Total | Survival | Total | Survival | Total | Survival | Total | Survival | Total animals | Average Survival | St. Dev. (+/-) | SEM (+/-) |
| 3 | 94 | 1.00 | 100 | 1.00 | 110 | 1.00 | 105 | 1.00 | 409 | 1.0000 | 0.0000 | 0.0000 |
| 5 | 83 | 1.00 | 88 | 1.00 | 158 | 1.00 | 99 | 1.00 | 428 | 1.0000 | 0.0000 | 0.0000 |
| 7 | 83 | 0.99 | 99 | 1.00 | 128 | 0.99 | 114 | 1.00 | 424 | 0.9950 | 0.0060 | 0.0030 |
| 9 | 99 | 1.00 | 91 | 1.00 | 103 | 0.99 | 89 | 0.99 | 382 | 0.9948 | 0.0061 | 0.0030 |
| 11 | 117 | 1.00 | 104 | 0.94 | 116 | 0.98 | 105 | 0.99 | 442 | 0.9789 | 0.0254 | 0.0127 |
| 13 | 94 | 0.88 | 111 | 0.99 | 125 | 0.98 | 107 | 0.74 | 437 | 0.8991 | 0.1180 | 0.0590 |
| 15 | 99 | 0.62 | 98 | 0.78 | 133 | 0.21 | 101 | 0.61 | 431 | 0.5540 | 0.2412 | 0.1206 |
| 17 | 90 | 0.63 | 87 | 0.74 | 143 | 0.06 | 132 | 0.24 | 452 | 0.4168 | 0.3210 | 0.1605 |
| 19 | 96 | 0.09 | 95 | 0.25 | 128 | 0.00 | 73 | 0.00 | 392 | 0.0866 | 0.1192 | 0.0596 |
| 21 | 100 | 0.16 | 100 | 0.38 | 110 | 0.00 | 102 | 0.00 | 412 | 0.1350 | 0.1799 | 0.0900 |
| ***age-1(hx546); ets-4(rrr16)*** | | | | | | | | | | | | |
| Days at 4°C | Total | Survival | Total | Survival | Total | Survival | Total | Survival | Total animals | Average Survival | St. Dev. (+/-) | SEM (+/-) |
| 3 | 105 | 1.00 | 85 | 1.00 | 107 | 1.00 | 127 | 1.00 | 424 | 1.0000 | 0.0000 | 0.0000 |
| 5 | 83 | 1.00 | 104 | 1.00 | 134 | 1.00 | 134 | 1.00 | 455 | 1.0000 | 0.0000 | 0.0000 |
| 7 | 85 | 1.00 | 110 | 1.00 | 125 | 1.00 | 93 | 1.00 | 413 | 1.0000 | 0.0000 | 0.0000 |
| 9 | 101 | 1.00 | 106 | 1.00 | 100 | 0.99 | 113 | 1.00 | 420 | 0.9975 | 0.0050 | 0.0025 |
| 11 | 95 | 0.97 | 88 | 0.93 | 114 | 0.85 | 164 | 0.99 | 461 | 0.9347 | 0.0605 | 0.0303 |
| 13 | 91 | 0.96 | 91 | 0.99 | 107 | 0.75 | 125 | 0.91 | 414 | 0.9012 | 0.1071 | 0.0535 |
| 15 | 107 | 0.90 | 112 | 0.73 | 129 | 0.58 | 147 | 0.49 | 495 | 0.6751 | 0.1786 | 0.0893 |
| 17 | 90 | 0.63 | 102 | 0.51 | 132 | 0.02 | 128 | 0.09 | 452 | 0.3111 | 0.3064 | 0.1532 |
| 19 | 94 | 0.09 | 78 | 0.55 | 109 | 0.14 | 102 | 0.13 | 383 | 0.2254 | 0.2185 | 0.1092 |
| 21 | 110 | 0.04 | 79 | 0.35 | 103 | 0.09 | 97 | 0.04 | 389 | 0.1299 | 0.1515 | 0.0757 |

Wilcoxon signed rank test

| | *age-1(hx546)* | *age-1(hx546); ets-4(rrr16)* | *ets-4(rrr16)* |
|---|---|---|---|
| *age-1(hx546);ets-4(rrr16)* | 0.6701 | | |
| *ets-4(rrr16)* | 0.0283 | 0.0042 | |
| wt | 5.4e-06 | 1.7e-06 | 1.7e-06 |

Note: the wt and *ets-4(rrr16)* replicates 1 and 2 were also used in 1E, and the replicates 3 and 4 in 4A (the additional strains were examined simultaneously)

| | 1st replicate | | 2nd replicate | | 3rd replicate | | 4th replicate | | | | | |
|---|---|---|---|---|---|---|---|---|---|---|---|---|
| **wt** | | | | | | | | | | | | |
| Days at 4°C | Total | Survival | Total | Survival | Total | Survival | Total | Survival | Total animals | Average Survival | St. Dev. (+/-) | SEM (+/-) |
| 3 | 108 | 0.99 | 105 | 1.00 | 94 | 1.00 | 112 | 1.00 | 419 | 0.9977 | 0.0046 | 0.0023 |
| 5 | 103 | 0.97 | 106 | 1.00 | 92 | 1.00 | 114 | 1.00 | 415 | 0.9927 | 0.0146 | 0.0073 |
| 7 | 116 | 0.97 | 97 | 1.00 | 100 | 0.91 | 112 | 0.22 | 425 | 0.7747 | 0.3695 | 0.1848 |
| 9 | 103 | 0.59 | 124 | 0.11 | 82 | 0.04 | 111 | 0.00 | 420 | 0.1854 | 0.2753 | 0.1376 |
| 11 | 102 | 0.04 | 96 | 0.10 | 74 | 0.00 | 115 | 0.00 | 387 | 0.0358 | 0.0492 | 0.0246 |
| 13 | 105 | 0.00 | 100 | 0.08 | 113 | 0.00 | 120 | 0.00 | 438 | 0.0200 | 0.0400 | 0.0200 |
| 15 | 94 | 0.01 | 146 | 0.00 | 89 | 0.00 | 141 | 0.01 | 470 | 0.0044 | 0.0053 | 0.0027 |
| ***ets-4(rrr16)*** | | | | | | | | | | | | |
| Days at 4°C | Total | Survival | Total | Survival | Total | Survival | Total | Survival | Total animals | Average Survival | St. Dev. (+/-) | SEM (+/-) |
| 3 | 95 | 1.00 | 92 | 1.00 | 139 | 1.00 | 137 | 1.00 | 463 | 1.0000 | 0.0000 | 0.0000 |
| 5 | 80 | 1.00 | 76 | 1.00 | 147 | 1.00 | 107 | 1.00 | 410 | 1.0000 | 0.0000 | 0.0000 |
| 7 | 75 | 0.99 | 77 | 0.99 | 157 | 0.99 | 154 | 0.74 | 463 | 0.9269 | 0.1245 | 0.0622 |
| 9 | 107 | 0.90 | 84 | 0.82 | 147 | 0.86 | 114 | 0.54 | 452 | 0.7816 | 0.1615 | 0.0808 |
| 11 | 80 | 0.71 | 94 | 0.47 | 129 | 0.77 | 116 | 0.30 | 419 | 0.5623 | 0.2173 | 0.1087 |
| 13 | 74 | 0.27 | 81 | 0.52 | 175 | 0.58 | 170 | 0.16 | 500 | 0.3812 | 0.1992 | 0.0996 |
| 15 | 62 | 0.29 | 83 | 0.40 | 136 | 0.20 | 162 | 0.10 | 443 | 0.2463 | 0.1276 | 0.0638 |
| ***daf-16(mu86); ets-4(rrr16)*** | | | | | | | | | | | | |
| Days at 4°C | Total | Survival | Total | Survival | Total | Survival | Total | Survival | Total animals | Average Survival | St. Dev. (+/-) | SEM (+/-) |
| 3 | 88 | 1.00 | 84 | 1.00 | 115 | 1.00 | 143 | 0.99 | 430 | 0.9983 | 0.0035 | 0.0017 |
| 5 | 68 | 1.00 | 85 | 0.98 | 120 | 0.99 | 78 | 0.90 | 351 | 0.9664 | 0.0470 | 0.0235 |
| 7 | 99 | 0.97 | 69 | 1.00 | 132 | 0.42 | 104 | 0.15 | 404 | 0.6369 | 0.4168 | 0.2084 |
| 9 | 97 | 0.69 | 79 | 0.63 | 117 | 0.01 | 85 | 0.02 | 378 | 0.3389 | 0.3736 | 0.1868 |
| 11 | 69 | 0.01 | 67 | 0.01 | 101 | 0.00 | 126 | 0.00 | 363 | 0.0074 | 0.0085 | 0.0042 |
| 13 | 97 | 0.03 | 79 | 0.01 | 113 | 0.00 | 91 | 0.00 | 380 | 0.0109 | 0.0146 | 0.0073 |
| 15 | 78 | 0.14 | 83 | 0.01 | 122 | 0.00 | 116 | 0.00 | 399 | 0.0383 | 0.0687 | 0.0344 |

Wilcoxon signed rank test

| | *daf-16(mu86); ets-4(rrr16)* | *ets-4(rrr16)* |
|---|---|---|
| *ets-4(rrr16)* | 7.5e-05 | |
| wt | 0.64 | 7.9e-05 |

Note: the wt and *ets-4(rrr16)* replicate 3 was also used in 4A (the additional strains were examined simultaneously)

| | 1st replicate | | 2nd replicate | | 3rd replicate | | 4th replicate | | | | | |
|---|---|---|---|---|---|---|---|---|---|---|---|---|
| **wt** | | | | | | | | | | | | |
| Days at 4°C | Total | Survival | Total | Survival | Total | Survival | Total | Survival | Total animals | Average Survival | St. Dev. (+/-) | SEM (+/-) |
| 3 | 149 | 0.99 | 162 | 1.00 | 106 | 0.95 | 161 | 1.00 | 578 | 0.9865 | 0.0227 | 0.0113 |
| 5 | 227 | 1.00 | 150 | 1.00 | 105 | 0.96 | 141 | 1.00 | 623 | 0.9905 | 0.0190 | 0.0095 |
| 7 | 137 | 0.55 | 172 | 0.62 | 135 | 0.19 | 119 | 0.51 | 563 | 0.4672 | 0.1881 | 0.0940 |
| 9 | 170 | 0.00 | 155 | 0.01 | 136 | 0.18 | 141 | 0.01 | 602 | 0.0493 | 0.0897 | 0.0449 |
| 11 | 146 | 0.03 | 96 | 0.00 | 115 | 0.00 | 104 | 0.00 | 461 | 0.0086 | 0.0171 | 0.0086 |
| 13 | 162 | 0.01 | 162 | 0.00 | 117 | 0.00 | 137 | 0.00 | 578 | 0.0015 | 0.0031 | 0.0015 |
| 15 | 126 | 0.00 | 149 | 0.00 | 121 | 0.00 | 108 | 0.00 | 504 | 0.0000 | 0.0000 | 0.0000 |
| ***ets-4(rrr16)*** | | | | | | | | | | | | |
| Days at 4°C | Total | Survival | Total | Survival | Total | Survival | Total | Survival | Total animals | Average Survival | St. Dev. (+/-) | SEM (+/-) |
| 3 | 135 | 1.00 | 120 | 1.00 | 140 | 1.00 | 110 | 0.98 | 505 | 0.9955 | 0.0091 | 0.0045 |
| 5 | 130 | 1.00 | 139 | 1.00 | 117 | 1.00 | 95 | 1.00 | 481 | 1.0000 | 0.0000 | 0.0000 |
| 7 | 181 | 0.98 | 156 | 0.99 | 115 | 0.96 | 101 | 0.89 | 553 | 0.9548 | 0.0451 | 0.0225 |
| 9 | 153 | 0.98 | 176 | 0.73 | 134 | 0.94 | 126 | 0.63 | 589 | 0.8207 | 0.1664 | 0.0832 |
| 11 | 168 | 0.74 | 193 | 0.32 | 151 | 0.61 | 128 | 0.45 | 640 | 0.5300 | 0.1852 | 0.0926 |
| 13 | 176 | 0.53 | 177 | 0.30 | 181 | 0.51 | 108 | 0.33 | 642 | 0.4187 | 0.1192 | 0.0596 |
| 15 | 177 | 0.45 | 147 | 0.26 | 175 | 0.54 | 110 | 0.13 | 609 | 0.3437 | 0.1865 | 0.0932 |
| ***pqm-1(ok485); ets-4(rrr16)*** | | | | | | | | | | | | |
| Days at 4°C | Total | Survival | Total | Survival | Total | Survival | Total | Survival | Total animals | Average Survival | St. Dev. (+/-) | SEM (+/-) |
| 3 | 139 | 0.99 | 148 | 1.00 | 105 | 1.00 | 94 | 1.00 | 486 | 0.9982 | 0.0036 | 0.0018 |
| 5 | 136 | 1.00 | 79 | 0.96 | 102 | 1.00 | 86 | 1.00 | 403 | 0.9905 | 0.0190 | 0.0095 |
| 7 | 109 | 0.61 | 98 | 0.92 | 110 | 0.22 | 99 | 0.81 | 416 | 0.6398 | 0.3078 | 0.1539 |
| 9 | 148 | 0.09 | 125 | 0.18 | 92 | 0.33 | 120 | 0.00 | 485 | 0.1492 | 0.1381 | 0.0691 |
| 11 | 126 | 0.02 | 103 | 0.00 | 112 | 0.00 | 92 | 0.03 | 433 | 0.0121 | 0.0156 | 0.0078 |
| 13 | 110 | 0.02 | 106 | 0.00 | 92 | 0.00 | 124 | 0.00 | 432 | 0.0045 | 0.0091 | 0.0045 |
| 15 | 102 | 0.00 | 74 | 0.00 | 112 | 0.00 | 98 | 0.01 | 386 | 0.0026 | 0.0051 | 0.0026 |
| ***daf-16(mu86); pqm-1(ok485); ets-4 (rrr16)*** | | | | | | | | | | | | |
| Days at 4°C | Total | Survival | Total | Survival | Total | Survival | Total | Survival | Total animals | Average Survival | St. Dev. (+/-) | SEM (+/-) |
| 3 | 171 | 0.99 | 152 | 1.00 | 92 | 0.99 | 139 | 0.97 | 554 | 0.9886 | 0.0124 | 0.0062 |
| 5 | 168 | 1.00 | 101 | 1.00 | 113 | 0.96 | 148 | 0.86 | 530 | 0.9574 | 0.0639 | 0.0319 |
| 7 | 123 | 0.67 | 151 | 0.12 | 127 | 0.01 | 166 | 0.16 | 567 | 0.2396 | 0.2969 | 0.1485 |
| 9 | 136 | 0.01 | 147 | 0.00 | 125 | 0.00 | 161 | 0.00 | 569 | 0.0037 | 0.0074 | 0.0037 |
| 11 | 144 | 0.01 | 154 | 0.00 | 96 | 0.00 | 103 | 0.01 | 497 | 0.0059 | 0.0070 | 0.0035 |
| 13 | 145 | 0.01 | 131 | 0.00 | 125 | 0.00 | 135 | 0.00 | 536 | 0.0017 | 0.0034 | 0.0017 |
| 15 | 103 | 0.00 | 128 | 0.00 | 97 | 0.00 | 145 | 0.00 | 473 | 0.0000 | 0.0000 | 0.0000 |

Wilcoxon signed rank test

| | *daf-16(mu86); pqm-1(ok485); ets-4 (rrr16)* | *ets-4(rrr16)* | *pqm-1(ok485); ets-4(rrr16)* |
|---|---|---|---|
| *ets-4(rrr16)* | 7.5e-05 | | |
| *pqm-1(ok485); ets-4(rrr16)* | 0.0100 | 7.5e-05 | |
| wt | 0.0796 | 7.5e-05 | 0.0095 |

| | 1st replicate | | 2nd replicate | | 3rd replicate | | | | | | | |
|---|---|---|---|---|---|---|---|---|---|---|---|---|
| **wt day 5 at 4°C** | | | | | | | | | | | **Treatment vs. control** | |
| RNAi condition | Total | Survival | Total | Survival | Total | Survival | Total animals | Average Survival | St. Dev. (+/-) | SEM (+/-) | p val | symbol |
| *EV* | 41 | 1.00 | 110 | 0.59 | 131 | 0.99 | 282 | 0.8609 | 0.2339 | 0.1350 | - | - |
| *asp-13* | 65 | 0.89 | 114 | 0.38 | 126 | 0.94 | 305 | 0.7380 | 0.3135 | 0.1810 | 0.1272 | n.s. |
| *ftn-1,-2* | 72 | 0.36 | 104 | 0.20 | 132 | 0.96 | 308 | 0.5084 | 0.4009 | 0.2315 | 0.1843 | n.s. |
| *fmo-2* | 45 | 0.87 | 104 | 0.67 | 131 | 0.98 | 280 | 0.8415 | 0.1573 | 0.0908 | 0.7853 | n.s. |
| *oac-14* | 53 | 0.72 | 109 | 0.80 | 126 | 0.48 | 288 | 0.6638 | 0.1675 | 0.0967 | 0.4525 | n.s. |
| *nhr-58* | 70 | 0.87 | 91 | 0.90 | 168 | 0.98 | 329 | 0.9162 | 0.0540 | 0.0312 | 0.7150 | n.s. |
| ***ets-4(rrr16)* day 5 at 4°C** | | | | | | | | | | | **Treatment vs. control** | |
| RNAi condition | Total | Survival | Total | Survival | Total | Survival | Total animals | Average Survival | St. Dev. (+/-) | SEM (+/-) | p val | symbol |
| *EV* | 53 | 0.98 | 98 | 0.95 | 91 | 0.92 | 242 | 0.9511 | 0.0291 | 0.0168 | - | - |
| *asp-13* | 39 | 0.67 | 108 | 0.93 | 97 | 0.99 | 244 | 0.8608 | 0.1711 | 0.0988 | 0.5147 | n.s. |
| *ftn-1,-2* | 58 | 0.45 | 120 | 0.34 | 108 | 0.88 | 286 | 0.5565 | 0.2848 | 0.1645 | 0.1554 | n.s. |
| *fmo-2* | 37 | 0.81 | 107 | 0.82 | 92 | 0.99 | 236 | 0.8741 | 0.0998 | 0.0576 | 0.4003 | n.s. |
| *oac-14* | 52 | 0.65 | 100 | 0.81 | 115 | 1.00 | 267 | 0.8213 | 0.1734 | 0.1001 | 0.3821 | n.s. |
| *nhr-58* | 50 | 1.00 | 91 | 0.73 | 94 | 0.96 | 235 | 0.8942 | 0.1479 | 0.0854 | 0.5667 | n.s. |
| **wt day 7 at 4°C** | | | | | | | | | | | **Treatment vs. control** | |
| RNAi condition | Total | Survival | Total | Survival | Total | Survival | Total animals | Average Survival | St. Dev. (+/-) | SEM (+/-) | p val | symbol |
| *EV* | 93 | 0.12 | 121 | 0.05 | 93 | 0.11 | 307 | 0.0918 | 0.0369 | 0.0213 | - | - |
| *asp-13* | 124 | 0.28 | 98 | 0.11 | 105 | 0.12 | 327 | 0.1728 | 0.0950 | 0.0548 | 0.2045 | n.s. |
| *ftn-1,-2* | 91 | 0.00 | 130 | 0.11 | 93 | 0.03 | 314 | 0.0467 | 0.0553 | 0.0319 | 0.4847 | n.s. |
| *fmo-2* | 143 | 0.52 | 134 | 0.07 | 112 | 0.38 | 389 | 0.3199 | 0.2302 | 0.1329 | 0.1784 | n.s. |
| *oac-14* | 100 | 0.26 | 136 | 0.24 | 106 | 0.20 | 342 | 0.2336 | 0.0319 | 0.0184 | 0.0409 | * |
| *nhr-58* | 91 | 0.25 | 161 | 0.06 | 97 | 0.01 | 349 | 0.1063 | 0.1288 | 0.0744 | 0.8485 | n.s. |
| ***ets-4(rrr16)* day 7 at 4°C** | | | | | | | | | | | **Treatment vs. control** | |
| RNAi condition | Total | Survival | Total | Survival | Total | Survival | Total animals | Average Survival | St. Dev. (+/-) | SEM (+/-) | p val | symbol |
| *EV* | 73 | 0.77 | 113 | 0.58 | 96 | 0.83 | 282 | 0.7252 | 0.1341 | 0.0774 | - | - |
| *asp-13* | 77 | 0.87 | 112 | 0.43 | 101 | 0.85 | 290 | 0.7167 | 0.2497 | 0.1442 | 0.9183 | n.s. |
| *ftn-1,-2* | 69 | 0.46 | 98 | 0.31 | 89 | 0.49 | 256 | 0.4214 | 0.1010 | 0.0583 | 0.0044 | * |
| *fmo-2* | 74 | 0.82 | 72 | 0.94 | 86 | 0.83 | 232 | 0.8648 | 0.0690 | 0.0398 | 0.3532 | n.s. |
| *oac-14* | 72 | 0.92 | 113 | 0.58 | 107 | 0.78 | 292 | 0.7588 | 0.1669 | 0.0964 | 0.6376 | n.s. |
| *nhr-58* | 75 | 0.93 | 117 | 0.59 | 93 | 0.80 | 285 | 0.7729 | 0.1729 | 0.0998 | 0.5170 | n.s. |

Two tailed, paired, student t.test:

| | |
|---|---|
| $p < 0.05$ | * |
| $p < 0.01$ | ** |
| $p < 0.001$ | *** |

| | 1st replicate | | 2nd replicate | | 3rd replicate | | | | | |
|---|---|---|---|---|---|---|---|---|---|---|
| **wt** | | | | | | | | | | |
| Days at 4°C | Total | Survival | Total | Survival | Total | Survival | Total animals | Average Survival | St. Dev. (+/-) | SEM (+/-) |
| 3 | 94 | 1.00 | 92 | 1.00 | 86 | 1.00 | 272 | 1.0000 | 0.0000 | 0.0000 |
| 5 | 92 | 1.00 | 94 | 1.00 | 150 | 1.00 | 336 | 1.0000 | 0.0000 | 0.0000 |
| 7 | 100 | 0.91 | 120 | 0.70 | 137 | 0.81 | 357 | 0.8067 | 0.1050 | 0.0606 |
| 9 | 82 | 0.04 | 121 | 0.02 | 112 | 0.02 | 315 | 0.0237 | 0.0112 | 0.0065 |
| 11 | 74 | 0.00 | 98 | 0.00 | 149 | 0.00 | 321 | 0.0000 | 0.0000 | 0.0000 |
| 13 | 113 | 0.00 | 154 | 0.00 | 129 | 0.00 | 396 | 0.0000 | 0.0000 | 0.0000 |
| 15 | 89 | 0.00 | 128 | 0.00 | 155 | 0.00 | 372 | 0.0000 | 0.0000 | 0.0000 |
| ***ets-4(rrr16)*** | | | | | | | | | | |
| Days at 4°C | Total | Survival | Total | Survival | Total | Survival | Total | Average Survival | St. Dev. (+/-) | SEM (+/-) |
| 3 | 139 | 1.00 | 101 | 1.00 | 97 | 0.99 | 337 | 0.9966 | 0.0060 | 0.0034 |
| 5 | 147 | 1.00 | 79 | 1.00 | 147 | 1.00 | 373 | 1.0000 | 0.0000 | 0.0000 |
| 7 | 157 | 0.99 | 68 | 1.00 | 154 | 0.97 | 379 | 0.9892 | 0.0135 | 0.0078 |
| 9 | 147 | 0.86 | 99 | 0.92 | 141 | 0.63 | 387 | 0.8048 | 0.1528 | 0.0882 |
| 11 | 129 | 0.77 | 66 | 0.91 | 181 | 0.52 | 376 | 0.7320 | 0.1973 | 0.1139 |
| 13 | 175 | 0.58 | 96 | 0.63 | 150 | 0.33 | 421 | 0.5118 | 0.1564 | 0.0903 |
| 15 | 136 | 0.20 | 91 | 0.51 | 135 | 0.24 | 362 | 0.3137 | 0.1672 | 0.0965 |
| ***ftn-1(ok485)*** | | | | | | | | | | |
| Days at 4°C | Total | Survival | Total | Survival | Total | Survival | Total animals | Average Survival | St. Dev. (+/-) | SEM (+/-) |
| 3 | 101 | 1.00 | 107 | 1.00 | 133 | 1.00 | 341 | 1.0000 | 0.0000 | 0.0000 |
| 5 | 121 | 0.99 | 97 | 0.98 | 133 | 1.00 | 351 | 0.9904 | 0.0104 | 0.0060 |
| 7 | 84 | 0.85 | 158 | 0.92 | 131 | 0.92 | 373 | 0.8955 | 0.0437 | 0.0252 |
| 9 | 113 | 0.01 | 134 | 0.17 | 104 | 0.08 | 351 | 0.0858 | 0.0818 | 0.0472 |
| 11 | 111 | 0.00 | 109 | 0.00 | 134 | 0.00 | 354 | 0.0000 | 0.0000 | 0.0000 |
| 13 | 120 | 0.00 | 110 | 0.00 | 135 | 0.00 | 365 | 0.0000 | 0.0000 | 0.0000 |
| 15 | 119 | 0.00 | 96 | 0.00 | 179 | 0.00 | 394 | 0.0000 | 0.0000 | 0.0000 |
| ***ftn-1(ok485); ets-4(rrr16)*** | | | | | | | | | | |
| Days at 4°C | Total | Survival | Total | Survival | Total | Survival | Total animals | Average Survival | St. Dev. (+/-) | SEM (+/-) |
| 3 | 50 | 1.00 | 97 | 1.00 | 92 | 1.00 | 239 | 1.0000 | 0.0000 | 0.0000 |
| 5 | 56 | 1.00 | 116 | 1.00 | 153 | 1.00 | 325 | 1.0000 | 0.0000 | 0.0000 |
| 7 | 68 | 0.50 | 134 | 0.90 | 161 | 0.83 | 363 | 0.7430 | 0.2139 | 0.1235 |
| 9 | 47 | 0.13 | 80 | 0.35 | 144 | 0.63 | 271 | 0.3699 | 0.2527 | 0.1459 |
| 11 | 46 | 0.02 | 119 | 0.05 | 137 | 0.00 | 302 | 0.0241 | 0.0253 | 0.0146 |
| 13 | 51 | 0.00 | 108 | 0.00 | 147 | 0.00 | 306 | 0.0000 | 0.0000 | 0.0000 |
| 15 | 42 | 0.00 | 93 | 0.01 | 165 | 0.00 | 300 | 0.0036 | 0.0062 | 0.0036 |

Wilcoxon signed rank test

| | *ets-4(rrr16)* | *ftn-1(ok485)* | *ftn-1(ok485); ets-4 (rrr16)* |
|---|---|---|---|
| *ftn-1(ok485)* | 0.0017 | | |
| *ftn-1(ok485); ets-4 (rrr16)* | 0.0018 | 0.3627 | |
| wt | 0.0018 | 0.3627 | 0.1458 |

| | 1st replicate | | 2nd replicate | | 3rd replicate | | | | | |
|---|---|---|---|---|---|---|---|---|---|---|
| **wt** | | | | | | | | | | |
| Days at 4°C | Total | Survival | Total | Survival | Total | Survival | Total animals | Avarage survival | St. Dev. (+/-) | SEM (+/-) |
| 3 | 112 | 0.98 | 79 | 0.97 | 86 | 1.00 | 277 | 0.9856 | 0.0130 | 0.0075 |
| 5 | 136 | 0.78 | 77 | 0.73 | 97 | 0.97 | 310 | 0.8253 | 0.1273 | 0.0735 |
| 7 | 126 | 0.28 | 88 | 0.08 | 84 | 0.33 | 298 | 0.2302 | 0.1334 | 0.0770 |
| 9 | 127 | 0.17 | 83 | 0.02 | 58 | 0.02 | 268 | 0.0715 | 0.0881 | 0.0509 |
| 11 | 108 | 0.00 | 71 | 0.00 | 82 | 0.00 | 261 | 0.0000 | 0.0000 | 0.0000 |
| 13 | 132 | 0.00 | 73 | 0.00 | 76 | 0.00 | 281 | 0.0000 | 0.0000 | 0.0000 |
| 15 | 116 | 0.00 | 90 | 0.00 | 74 | 0.00 | 280 | 0.0000 | 0.0000 | 0.0000 |
| ***ftn-1(syb2550)*** | | | | | | | | | | |
| Days at 4°C | Total | Survival | Total | Survival | Total | Survival | Total animals | Avarage survival | St. Dev. (+/-) | SEM (+/-) |
| 3 | 153 | 0.99 | 147 | 0.99 | 96 | 0.98 | 396 | 0.9886 | 0.0082 | 0.0047 |
| 5 | 114 | 0.50 | 123 | 0.65 | 113 | 0.88 | 350 | 0.6755 | 0.1893 | 0.1093 |
| 7 | 113 | 0.14 | 169 | 0.04 | 87 | 0.13 | 369 | 0.1031 | 0.0540 | 0.0312 |
| 9 | 97 | 0.08 | 159 | 0.01 | 94 | 0.01 | 350 | 0.0331 | 0.0428 | 0.0247 |
| 11 | 130 | 0.00 | 117 | 0.00 | 104 | 0.00 | 351 | 0.0000 | 0.0000 | 0.0000 |
| 13 | 101 | 0.00 | 135 | 0.00 | 86 | 0.00 | 322 | 0.0000 | 0.0000 | 0.0000 |
| 15 | 95 | 0.00 | 110 | 0.00 | 103 | 0.00 | 308 | 0.0000 | 0.0000 | 0.0000 |
| ***ets-4(rrr16)*** | | | | | | | | | | |
| Days at 4°C | Total | Survival | Total | Survival | Total | Survival | Total animals | Avarage survival | St. Dev. (+/-) | SEM (+/-) |
| 3 | 132 | 1.00 | 127 | 0.99 | 102 | 0.98 | 361 | 0.9908 | 0.0099 | 0.0057 |
| 5 | 115 | 0.92 | 107 | 0.98 | 90 | 0.98 | 312 | 0.9603 | 0.0334 | 0.0193 |
| 7 | 127 | 0.91 | 96 | 0.52 | 101 | 0.89 | 324 | 0.7751 | 0.2205 | 0.1273 |
| 9 | 96 | 0.91 | 103 | 0.37 | 103 | 0.55 | 302 | 0.6095 | 0.2730 | 0.1576 |
| 11 | 102 | 0.25 | 93 | 0.37 | 95 | 0.40 | 290 | 0.3369 | 0.0813 | 0.0470 |
| 13 | 97 | 0.31 | 89 | 0.15 | 99 | 0.33 | 285 | 0.2629 | 0.1019 | 0.0588 |
| 15 | 172 | 0.17 | 98 | 0.17 | 86 | 0.15 | 356 | 0.1664 | 0.0132 | 0.0076 |
| ***ftn-1(syb2550); ets-4(rrr16)*** | | | | | | | | | | |
| Days at 4°C | Total | Survival | Total | Survival | Total | Survival | Total animals | Avarage survival | St. Dev. (+/-) | SEM (+/-) |
| 3 | 98 | 0.95 | 74 | 0.93 | 130 | 0.96 | 302 | 0.9477 | 0.0146 | 0.0084 |
| 5 | 102 | 0.32 | 81 | 0.48 | 134 | 0.85 | 317 | 0.5519 | 0.2706 | 0.1562 |
| 7 | 113 | 0.19 | 94 | 0.10 | 130 | 0.30 | 337 | 0.1939 | 0.1024 | 0.0591 |
| 9 | 81 | 0.05 | 71 | 0.00 | 129 | 0.00 | 281 | 0.0165 | 0.0285 | 0.0165 |
| 11 | 85 | 0.00 | 96 | 0.00 | 134 | 0.00 | 315 | 0.0000 | 0.0000 | 0.0000 |
| 13 | 116 | 0.01 | 99 | 0.00 | 147 | 0.01 | 362 | 0.0051 | 0.0045 | 0.0026 |
| 15 | 104 | 0.00 | 83 | 0.00 | 126 | 0.00 | 313 | 0.0000 | 0.0000 | 0.0000 |

Wilcoxon signed rank test

| | *ets-4(rrr16)* | *ftn-1(syb2550)* | *ftn-1(syb2550); ets-4 (rrr16)* |
|---|---|---|---|
| *ftn-1(syb2550)* | 0.00023 | | |
| *ftn-1(syb2550); ets-4(rrr16)* | 0.00023 | 0.31517 | |
| wt | 0.00023 | 0.01294 | 0.00582 |

| | 1st replicate | | 2nd replicate | | 3rd replicate | | | | | |
|---|---|---|---|---|---|---|---|---|---|---|
| **wt** | | | | | | | | | | |
| Days at 4°C | Total | Survival | Total | Survival | Total | Survival | Total animals | Avarage survival | St. Dev. (+/-) | SEM (+/-) |
| 3 | 94 | 1.00 | 105 | 1.00 | 50 | 0.96 | 249 | 0.99 | 0.02 | 0.01 |
| 5 | 100 | 0.93 | 85 | 0.99 | 88 | 0.98 | 273 | 0.97 | 0.03 | 0.02 |
| 7 | 61 | 0.03 | 105 | 0.73 | 112 | 0.21 | 278 | 0.32 | 0.36 | 0.21 |
| 9 | 98 | 0.01 | 97 | 0.09 | 70 | 0.01 | 265 | 0.04 | 0.05 | 0.03 |
| 11 | 92 | 0.00 | 91 | 0.00 | 93 | 0.01 | 276 | 0.00 | 0.01 | 0.00 |
| 13 | 127 | 0.00 | 99 | 0.00 | 81 | 0.00 | 307 | 0.00 | 0.00 | 0.00 |
| ***sybSi67[Pdpy-30::ftn-1::unc-54 3'UTR];unc-119(ed3)*** | | | | | | | | | | |
| Days at 4°C | Total | Survival | Total | Survival | Total | Survival | Total animals | Avarage survival | St. Dev. (+/-) | SEM (+/-) |
| 3 | 103 | 1.00 | 83 | 1.00 | 67 | 1.00 | 253 | 1.00 | 0.00 | 0.00 |
| 5 | 93 | 0.98 | 83 | 0.99 | 59 | 0.98 | 235 | 0.98 | 0.00 | 0.00 |
| 7 | 100 | 0.79 | 67 | 1.00 | 91 | 0.99 | 258 | 0.93 | 0.12 | 0.07 |
| 9 | 85 | 0.20 | 93 | 0.86 | 55 | 0.62 | 233 | 0.56 | 0.33 | 0.19 |
| 11 | 80 | 0.06 | 68 | 0.22 | 122 | 0.13 | 270 | 0.14 | 0.08 | 0.05 |
| 13 | 115 | 0.01 | 47 | 0.00 | 83 | 0.01 | 245 | 0.01 | 0.01 | 0.00 |
| ***sybSi72[Pvit-5::ftn-1::unc-54 3'UTR];unc-119(ed3)*** | | | | | | | | | | |
| Days at 4°C | Total | Survival | Total | Survival | Total | Survival | Total animals | Avarage survival | St. Dev. (+/-) | SEM (+/-) |
| 3 | 92 | 1.00 | 87 | 1.00 | 53 | 1.00 | 232 | 1.00 | 0.00 | 0.00 |
| 5 | 109 | 0.97 | 78 | 1.00 | 74 | 0.97 | 261 | 0.98 | 0.02 | 0.01 |
| 7 | 142 | 0.83 | 91 | 0.98 | 71 | 1.00 | 304 | 0.94 | 0.09 | 0.05 |
| 9 | 95 | 0.49 | 100 | 0.76 | 71 | 0.85 | 266 | 0.70 | 0.18 | 0.11 |
| 11 | 117 | 0.11 | 86 | 0.19 | 89 | 0.70 | 292 | 0.33 | 0.32 | 0.18 |
| 13 | 94 | 0.00 | 85 | 0.04 | 71 | 0.24 | 250 | 0.09 | 0.13 | 0.07 |

**Wilcoxon signed rank test**

| | *sybSi67;unc-119(ed3)* | *sybSi72;unc-119(ed3)* |
|---|---|---|
| *sybSi72;unc-119(ed3)* | 0.0938 | |
| wt | 0.0013 | 0.0013 |

| | 1st replicate | | 2nd replicate | | 3rd replicate | | | | | |
|---|---|---|---|---|---|---|---|---|---|---|
| **wt** | | | | | | | | | | |
| Days at 4°C | Total | Survival | Total | Survival | Total | Survival | Total animals | Avarage survival | St. Dev. (+/-) | SEM (+/-) |
| 3 | 213 | 0.98 | 150 | 0.98 | 102 | 1.00 | 465 | 0.9855 | 0.0127 | 0.0073 |
| 5 | 177 | 0.75 | 197 | 0.62 | 136 | 0.96 | 510 | 0.7797 | 0.1712 | 0.0988 |
| 7 | 157 | 0.06 | 188 | 0.00 | 151 | 0.81 | 496 | 0.2884 | 0.4508 | 0.2603 |
| 9 | 196 | 0.01 | 262 | 0.00 | 167 | 0.34 | 625 | 0.1147 | 0.1910 | 0.1103 |
| 11 | 163 | 0.00 | 242 | 0.00 | 127 | 0.10 | 532 | 0.0341 | 0.0591 | 0.0341 |
| 13 | 164 | 0.00 | 169 | 0.00 | 167 | 0.05 | 500 | 0.0160 | 0.0277 | 0.0160 |
| 15 | 157 | 0.00 | 221 | 0.00 | 178 | 0.02 | 556 | 0.0075 | 0.0130 | 0.0075 |
| 17 | 159 | 0.00 | 305 | 0.00 | 154 | 0.00 | 618 | 0.0000 | 0.0000 | 0.0000 |
| 19 | 162 | 0.00 | 216 | 0.00 | 140 | 0.00 | 518 | 0.0000 | 0.0000 | 0.0000 |
| 21 | 135 | 0.00 | 184 | 0.00 | 113 | 0.00 | 432 | 0.0000 | 0.0000 | 0.0000 |
| 23 | 143 | 0.00 | 110 | 0.00 | 71 | 0.00 | 324 | 0.0000 | 0.0000 | 0.0000 |
| ***daf-2(e1370)*** | | | | | | | | | | |
| Days at 4°C | Total | Survival | Total | Survival | Total | Survival | Total animals | Avarage survival | St. Dev. (+/-) | SEM (+/-) |
| 3 | 183 | 1.00 | 172 | 1.00 | 132 | 1.00 | 487 | 1.0000 | 0.0000 | 0.0000 |
| 5 | 166 | 0.99 | 148 | 1.00 | 84 | 0.99 | 398 | 0.9920 | 0.0069 | 0.0040 |
| 7 | 155 | 0.97 | 210 | 0.98 | 126 | 0.99 | 491 | 0.9824 | 0.0090 | 0.0052 |
| 9 | 173 | 0.95 | 169 | 0.96 | 107 | 0.97 | 449 | 0.9634 | 0.0092 | 0.0053 |
| 11 | 161 | 0.91 | 184 | 0.91 | 116 | 0.95 | 461 | 0.9227 | 0.0224 | 0.0129 |
| 13 | 124 | 0.52 | 168 | 0.67 | 180 | 0.82 | 472 | 0.6692 | 0.1463 | 0.0844 |
| 15 | 146 | 0.08 | 215 | 0.30 | 197 | 0.48 | 558 | 0.2874 | 0.2002 | 0.1156 |
| 17 | 170 | 0.04 | 170 | 0.05 | 206 | 0.16 | 546 | 0.0828 | 0.0676 | 0.0390 |
| 19 | 176 | 0.01 | 183 | 0.01 | 142 | 0.02 | 501 | 0.0126 | 0.0079 | 0.0045 |
| 21 | 121 | 0.03 | 151 | 0.01 | 153 | 0.03 | 425 | 0.0241 | 0.0101 | 0.0058 |
| 23 | 143 | 0.00 | 189 | 0.01 | 159 | 0.00 | 491 | 0.0018 | 0.0031 | 0.0018 |
| ***ets-4(rrr16)*** | | | | | | | | | | |
| Days at 4°C | Total | Survival | Total | Survival | Total | Survival | Total animals | Avarage survival | St. Dev. (+/-) | SEM (+/-) |
| 3 | 201 | 0.99 | 198 | 0.99 | 190 | 1.00 | 589 | 0.9917 | 0.0076 | 0.0044 |
| 5 | 173 | 0.99 | 266 | 0.91 | 98 | 0.98 | 537 | 0.9599 | 0.0473 | 0.0273 |
| 7 | 209 | 0.73 | 174 | 0.44 | 134 | 0.99 | 517 | 0.7180 | 0.2744 | 0.1584 |
| 9 | 237 | 0.81 | 231 | 0.17 | 144 | 0.88 | 612 | 0.6189 | 0.3916 | 0.2261 |
| 11 | 160 | 0.31 | 227 | 0.11 | 141 | 0.59 | 528 | 0.3371 | 0.2402 | 0.1387 |
| 13 | 203 | 0.22 | 195 | 0.06 | 130 | 0.36 | 528 | 0.2149 | 0.1501 | 0.0867 |
| 15 | 177 | 0.16 | 321 | 0.04 | 170 | 0.18 | 668 | 0.1240 | 0.0756 | 0.0436 |
| 17 | 197 | 0.09 | 207 | 0.02 | 102 | 0.25 | 506 | 0.1169 | 0.1160 | 0.0669 |
| 19 | 147 | 0.09 | 248 | 0.01 | 160 | 0.13 | 555 | 0.0759 | 0.0625 | 0.0361 |
| 21 | 185 | 0.08 | 232 | 0.00 | 119 | 0.07 | 536 | 0.0476 | 0.0415 | 0.0239 |
| 23 | 173 | 0.00 | 196 | 0.00 | 131 | 0.05 | 500 | 0.0178 | 0.0309 | 0.0178 |
| ***daf-2(e1370); ets-4(rrr16)*** | | | | | | | | | | |
| Days at 4°C | Total | Survival | Total | Survival | Total | Survival | Total animals | Avarage survival | St. Dev. (+/-) | SEM (+/-) |
| 3 | 66 | 1.00 | 145 | 1.00 | 215 | 1.00 | 426 | 1.0000 | 0.0000 | 0.0000 |
| 5 | 95 | 0.99 | 124 | 0.99 | 217 | 1.00 | 436 | 0.9938 | 0.0055 | 0.0032 |
| 7 | 63 | 0.98 | 113 | 0.98 | 197 | 1.00 | 373 | 0.9888 | 0.0097 | 0.0056 |
| 9 | 50 | 1.00 | 115 | 0.97 | 163 | 0.99 | 328 | 0.9872 | 0.0131 | 0.0075 |
| 11 | 100 | 0.92 | 110 | 0.98 | 221 | 0.98 | 431 | 0.9597 | 0.0345 | 0.0199 |
| 13 | 97 | 0.84 | 115 | 0.90 | 166 | 0.85 | 378 | 0.8629 | 0.0366 | 0.0211 |
| 15 | 71 | 0.21 | 91 | 0.68 | 228 | 0.70 | 390 | 0.5314 | 0.2775 | 0.1602 |
| 17 | 70 | 0.13 | 94 | 0.31 | 211 | 0.38 | 375 | 0.2721 | 0.1292 | 0.0746 |
| 19 | 68 | 0.00 | 114 | 0.14 | 222 | 0.27 | 404 | 0.1354 | 0.1330 | 0.0768 |
| 21 | 81 | 0.06 | 95 | 0.13 | 225 | 0.20 | 401 | 0.1293 | 0.0692 | 0.0399 |
| 23 | 76 | 0.03 | 86 | 0.09 | 198 | 0.19 | 360 | 0.1021 | 0.0807 | 0.0466 |

Wilcoxon signed rank test

| | *daf-2(e1370)* | *daf-2(e1370); ets-4(rrr16)* | *ets-4(rrr16)* |
|---|---|---|---|
| *daf-2(e1370); ets-4(rrr16* | 4.9e-06 | | |
| *ets-4(rrr16)* | 0.0084 | 4.9e-06 | |
| wt | 4.9e-06 | 4.9e-06 | 4.9e-06 |

| | 1st replicate | | 2nd replicate | | 3rd replicate | | | | | | | |
|---|---|---|---|---|---|---|---|---|---|---|---|---|
| **wt** | | | | | | | | | | | | |
| Days at 4°C | Total | Survival | Total | Survival | Total | Survival | Total animals | Average Survival | St. Dev. (+/-) | SEM (+/-) | | |
| 3 | 95 | 1.00 | 83 | 0.99 | 96 | 0.99 | 274 | 0.9925 | 0.0065 | 0.0038 | | |
| 5 | 75 | 1.00 | 45 | 1.00 | 70 | 1.00 | 190 | 1.0000 | 0.0000 | 0.0000 | | |
| 7 | 87 | 0.99 | 89 | 0.99 | 100 | 1.00 | 276 | 0.9924 | 0.0066 | 0.0038 | | |
| 9 | 94 | 0.34 | 66 | 0.17 | 100 | 0.09 | 260 | 0.1990 | 0.1283 | 0.0741 | | |
| 11 | 75 | 0.00 | 87 | 0.15 | 91 | 0.20 | 253 | 0.1157 | 0.1031 | 0.0595 | | |
| ***age-1(hx546)*** | | | | | | | | | | | **vs. wt** | |
| Days at 4°C | Total | Survival | Total | Survival | Total | Survival | Total | Average Survival | St. Dev. (+/-) | SEM (+/-) | p val | symbol |
| 3 | 61 | 1.00 | 112 | 1.00 | 149 | 1.00 | 322 | 1.0000 | 0.0000 | 0.0000 | 0.1856 | n.s. |
| 5 | 47 | 1.00 | 57 | 0.98 | 143 | 1.00 | 247 | 0.9942 | 0.0101 | 0.0058 | 0.4226 | n.s. |
| 7 | 42 | 1.00 | 82 | 1.00 | 123 | 1.00 | 247 | 1.0000 | 0.0000 | 0.0000 | 0.1836 | n.s. |
| 9 | 50 | 1.00 | 53 | 1.00 | 110 | 0.97 | 213 | 0.9909 | 0.0157 | 0.0091 | 0.0072 | ** |
| 11 | 63 | 1.00 | 84 | 0.85 | 112 | 1.00 | 259 | 0.9484 | 0.0894 | 0.0516 | 0.0113 | ** |
| ***daf-16(mu86); age-1(hx546)*** | | | | | | | | | | | **vs. wt** | |
| Days at 4°C | Total | Survival | Total | Survival | Total | Survival | Total animals | Average Survival | St. Dev. (+/-) | SEM (+/-) | p val | symbol |
| 3 | 76 | 1.00 | 105 | 0.98 | 159 | 1.00 | 340 | 0.9937 | 0.0110 | 0.0063 | 0.8428 | n.s. |
| 5 | 65 | 1.00 | 113 | 0.99 | 123 | 1.00 | 301 | 0.9971 | 0.0051 | 0.0029 | 0.4226 | n.s. |
| 7 | 79 | 0.99 | 133 | 0.86 | 169 | 0.89 | 381 | 0.9127 | 0.0672 | 0.0388 | 0.1841 | n.s. |
| 9 | 63 | 0.11 | 112 | 0.00 | 130 | 0.03 | 305 | 0.0473 | 0.0574 | 0.0331 | 0.0925 | n.s. |
| 11 | 60 | 0.00 | 127 | 0.01 | 182 | 0.06 | 369 | 0.0228 | 0.0329 | 0.0190 | 0.1836 | n.s. |

Two tailed, paired, student t.test:

| | |
|---|---|
| p < 0.05 | * |
| p < 0.01 | ** |
| p < 0.001 | *** |

Note: since we did not observe the *age-1* mutants long enough to register the loss of viability, the data is unsuitable for Wilcoxon signed rank test. Thus, we used student t-test to compare the survival at 9 and 11 days, as shown in S1D.

Note': the wt replicates 1, 2 and 3 were also used in S1E (the additional strains were examined simultaneously)

| | 1st replicate | | 2nd replicate | | 3rd replicate | | | | | |
|---|---|---|---|---|---|---|---|---|---|---|
| ***wt*** | | | | | | | | | | |
| Days at 4°C | Total | Survival | Total | Survival | Total | Survival | Total animals | Average Survival | St. Dev. (+/-) | SEM (+/-) |
| 3 | 95 | 1.00 | 83 | 0.99 | 96 | 0.99 | 274 | 0.9925 | 0.0065 | 0.0038 |
| 5 | 75 | 1.00 | 45 | 1.00 | 70 | 1.00 | 190 | 1.0000 | 0.0000 | 0.0000 |
| 7 | 87 | 0.99 | 89 | 0.99 | 100 | 1.00 | 276 | 0.9924 | 0.0066 | 0.0038 |
| 9 | 94 | 0.34 | 66 | 0.17 | 100 | 0.09 | 260 | 0.1990 | 0.1283 | 0.0741 |
| 11 | 75 | 0.00 | 87 | 0.15 | 91 | 0.20 | 253 | 0.1157 | 0.1031 | 0.0595 |
| ***daf-16(mu86)*** | | | | | | | | | | |
| Days at 4°C | Total | Survival | Total | Survival | Total | Survival | Total | Average Survival | St. Dev. (+/-) | SEM (+/-) |
| 3 | 90 | 1.00 | 88 | 0.99 | 147 | 0.99 | 325 | 0.9917 | 0.0073 | 0.0042 |
| 5 | 87 | 0.99 | 76 | 1.00 | 127 | 0.97 | 290 | 0.9857 | 0.0159 | 0.0092 |
| 7 | 74 | 0.97 | 98 | 0.80 | 111 | 0.90 | 283 | 0.8899 | 0.0890 | 0.0514 |
| 9 | 71 | 0.41 | 98 | 0.01 | 101 | 0.05 | 270 | 0.1561 | 0.2195 | 0.1267 |
| 11 | 109 | 0.00 | 103 | 0.00 | 85 | 0.01 | 297 | 0.0039 | 0.0068 | 0.0039 |
| ***pqm-1(ok485)*** | | | | | | | | | | |
| Days at 4°C | Total | Survival | Total | Survival | Total | Survival | Total animals | Average Survival | St. Dev. (+/-) | SEM (+/-) |
| 3 | 101 | 1.00 | 119 | 0.99 | 110 | 1.00 | 330 | 0.9972 | 0.0049 | 0.0028 |
| 5 | 87 | 0.90 | 97 | 1.00 | 94 | 0.98 | 278 | 0.9584 | 0.0546 | 0.0315 |
| 7 | 72 | 1.00 | 91 | 0.98 | 127 | 0.96 | 290 | 0.9796 | 0.0197 | 0.0114 |
| 9 | 75 | 0.93 | 89 | 0.03 | 62 | 0.11 | 226 | 0.3600 | 0.4981 | 0.2876 |
| 11 | 59 | 0.08 | 106 | 0.00 | 68 | 0.10 | 233 | 0.0626 | 0.0549 | 0.0317 |
| ***daf-16(mu86); pqm-1(ok485)*** | | | | | | | | | | |
| Days at 4°C | Total | Survival | Total | Survival | Total | Survival | Total animals | Average Survival | St. Dev. (+/-) | SEM (+/-) |
| 3 | 136 | 1.00 | 66 | 1.00 | 59 | 1.00 | 261 | 1.0000 | 0.0000 | 0.0000 |
| 5 | 112 | 1.00 | 74 | 0.99 | 36 | 0.97 | 222 | 0.9862 | 0.0139 | 0.0080 |
| 7 | 88 | 0.99 | 69 | 0.91 | 50 | 0.94 | 207 | 0.9472 | 0.0383 | 0.0221 |
| 9 | 109 | 0.28 | 80 | 0.00 | 51 | 0.02 | 240 | 0.1013 | 0.1588 | 0.0917 |
| 11 | 101 | 0.00 | 71 | 0.00 | 74 | 0.00 | 246 | 0.0000 | 0.0000 | 0.0000 |

Wilcoxon signed rank test

| | *daf-16(mu86)* | *daf-16(mu86); pqm-1(ok48* | *pqm-1(ok485)* |
|---|---|---|---|
| *daf-16(mu86); pqm-1(ok485* | 0.592 | | |
| *pqm-1(ok485)* | 0.059 | 0.061 | |
| wt | 0.059 | 0.059 | 0.439 |

| | 1st replicate | | 2nd replicate | | 3rd replicate | | | | | |
|---|---|---|---|---|---|---|---|---|---|---|
| **wt CRTL** | | | | | | | | | | |
| Days at 4°C | Total | Survival | Total | Survival | Total | Survival | Total animals | Average Survival | St. Dev. (+/-) | SEM (+/-) |
| 5 | 122 | 0.69 | 103 | 0.99 | 71 | 1.00 | 296 | 0.8929 | 0.1771 | 0.1022 |
| 7 | 97 | 0.02 | 102 | 0.97 | 97 | 0.15 | 296 | 0.3819 | 0.5142 | 0.2969 |
| 9 | 79 | 0.01 | 61 | 0.07 | 118 | 0.00 | 258 | 0.0261 | 0.0348 | 0.0201 |
| 11 | 103 | 0.00 | 79 | 0.04 | 93 | 0.00 | 275 | 0.0127 | 0.0219 | 0.0127 |
| **wt 2.5 mM FAC** | | | | | | | | | | |
| Days at 4°C | Total | Survival | Total | Survival | Total | Survival | Total | Average Survival | St. Dev. (+/-) | SEM (+/-) |
| 5 | 109 | 0.61 | 74 | 1.00 | 95 | 0.99 | 278 | 0.8650 | 0.2127 | 0.1228 |
| 7 | 193 | 0.01 | 82 | 0.73 | 70 | 0.23 | 345 | 0.3233 | 0.2972 | 0.1716 |
| 9 | 66 | 0.00 | 96 | 0.05 | 97 | 0.00 | 259 | 0.0174 | 0.0920 | 0.0531 |
| 11 | 118 | 0.00 | 87 | 0.00 | 86 | 0.00 | 291 | 0.0000 | 0.2113 | 0.1220 |
| **wt 5 mM FAC** | | | | | | | | | | |
| Days at 4°C | Total | Survival | Total | Survival | Total | Survival | Total animals | Average Survival | St. Dev. (+/-) | SEM (+/-) |
| 5 | 72 | 0.17 | 80 | 0.95 | 122 | 0.98 | 274 | 0.7001 | 0.1811 | 0.1045 |
| 7 | 84 | 0.14 | 85 | 0.42 | 93 | 0.03 | 262 | 0.1995 | 0.2247 | 0.1297 |
| 9 | 86 | 0.00 | 64 | 0.02 | 100 | 0.00 | 250 | 0.0052 | 0.1493 | 0.0862 |
| 11 | 107 | 0.00 | 74 | 0.00 | 99 | 0.00 | 280 | 0.0000 | 0.2113 | 0.1220 |
| **wt 10 mM FAC** | | | | | | | | | | |
| Days at 4°C | Total | Survival | Total | Survival | Total | Survival | Total animals | Average Survival | St. Dev. (+/-) | SEM (+/-) |
| 5 | 111 | 0.14 | 80 | 0.96 | 90 | 0.62 | 281 | 0.5763 | 0.0865 | 0.0499 |
| 7 | 114 | 0.14 | 95 | 0.41 | 127 | 0.06 | 336 | 0.2020 | 0.2319 | 0.1339 |
| 9 | 101 | 0.00 | 82 | 0.00 | 97 | 0.00 | 280 | 0.0000 | 0.1618 | 0.0934 |
| 11 | 122 | 0.00 | 97 | 0.00 | 112 | 0.00 | 331 | 0.0000 | 0.2113 | 0.1220 |
| **wt 20 mM FAC** | | | | | | | | | | |
| Days at 4°C | Total | Survival | Total | Survival | Total | Survival | Total animals | Average Survival | St. Dev. (+/-) | SEM (+/-) |
| 5 | 142 | 0.33 | 71 | 0.55 | 76 | 0.34 | 289 | 0.4075 | 0.0162 | 0.0093 |
| 7 | 89 | 0.01 | 73 | 0.00 | 66 | 0.12 | 228 | 0.0441 | 0.1988 | 0.1148 |
| 9 | 100 | 0.00 | 79 | 0.01 | 87 | 0.00 | 266 | 0.0042 | 0.1521 | 0.0878 |
| 11 | 86 | 0.00 | 77 | 0.00 | 85 | 0.00 | 248 | 0.0000 | 0.2113 | 0.1220 |
| **wt 40 mM FAC** | | | | | | | | | | |
| Days at 4°C | Total | Survival | Total | Survival | Total | Survival | Total animals | Average Survival | St. Dev. (+/-) | SEM (+/-) |
| 5 | 90 | 0.01 | 66 | 0.02 | 98 | 0.00 | 254 | 0.0088 | 0.0067 | 0.0039 |
| 7 | 106 | 0.00 | 58 | 0.16 | 116 | 0.00 | 280 | 0.0517 | 0.1555 | 0.0898 |
| 9 | 87 | 0.00 | 58 | 0.00 | 49 | 0.02 | 194 | 0.0068 | 0.2610 | 0.1507 |
| 11 | 76 | 0.00 | 59 | 0.00 | 67 | 0.00 | 202 | 0.0000 | 0.2113 | 0.1220 |

**Wilcoxon signed rank test**

| | wt 10 mM FAC | wt 20 mM FAC | wt 2.5 mM FAC | wt 40 mM FAC | wt 5 mM FAC |
|---|---|---|---|---|---|
| wt 20 mM FAC | 0.2719 | | | | |
| wt 2.5 mM FAC | 0.0759 | 0.0360 | | | |
| wt 40 mM FAC | 0.0346 | 0.1829 | 0.0300 | | |
| wt 5 mM FAC | 0.3991 | 0.2049 | 0.1083 | 0.0249 | |
| wt CRTL | 0.0580 | 0.0091 | 0.2127 | 0.0125 | 0.0756 |

| | 1st replicate | | 2nd replicate | | 3rd replicate | | | | | |
|---|---|---|---|---|---|---|---|---|---|---|
| **wt CRTL** | | | | | | | | | | |
| Days at 4°C | Total | Survival | Total | Survival | Total | Survival | Total animals | Average Survival | St. Dev. (+/-) | SEM (+/-) |
| 5 | 227 | 1.00 | 150 | 1.00 | 105 | 0.96 | 482 | 0.9873 | 0.0220 | 0.0127 |
| 7 | 173 | 0.43 | 172 | 0.62 | 129 | 0.95 | 474 | 0.6652 | 0.2596 | 0.1499 |
| 9 | 170 | 0.00 | 155 | 0.01 | 116 | 0.00 | 441 | 0.0022 | 0.0037 | 0.0022 |
| 11 | 146 | 0.03 | 96 | 0.00 | 131 | 0.00 | 373 | 0.0114 | 0.0198 | 0.0114 |
| **wt 30 mM FAC** | | | | | | | | | | |
| Days at 4°C | Total | Survival | Total | Survival | Total | Survival | Total animals | Average Survival | St. Dev. (+/-) | SEM (+/-) |
| 5 | 201 | 0.68 | 145 | 0.43 | 136 | 0.85 | 482 | 0.6563 | 0.2104 | 0.1215 |
| 7 | 170 | 0.16 | 191 | 0.00 | 109 | 0.03 | 470 | 0.0621 | 0.0849 | 0.0490 |
| 9 | 184 | 0.00 | 162 | 0.00 | 80 | 0.01 | 426 | 0.0042 | 0.0072 | 0.0042 |
| 11 | 166 | 0.01 | 169 | 0.00 | 110 | 0.00 | 445 | 0.0020 | 0.0035 | 0.0020 |
| ***ets-4(rrr16)* CTRL** | | | | | | | | | | |
| Days at 4°C | Total | Survival | Total | Survival | Total | Survival | Total animals | Average Survival | St. Dev. (+/-) | SEM (+/-) |
| 5 | 130 | 1.00 | 139 | 1.00 | 117 | 1.00 | 386 | 1.0000 | 0.0000 | 0.0000 |
| 7 | 181 | 0.98 | 156 | 0.99 | 113 | 1.00 | 450 | 0.9905 | 0.0114 | 0.0066 |
| 9 | 153 | 0.98 | 176 | 0.73 | 132 | 0.90 | 461 | 0.8697 | 0.1295 | 0.0748 |
| 11 | 168 | 0.74 | 193 | 0.32 | 139 | 0.89 | 500 | 0.6525 | 0.2962 | 0.1710 |
| ***ets-4(rrr16)* 30 mM FAC** | | | | | | | | | | |
| Days at 4°C | Total | Survival | Total | Survival | Total | Survival | Total animals | Average Survival | St. Dev. (+/-) | SEM (+/-) |
| 5 | 164 | 0.90 | 154 | 0.85 | 139 | 0.96 | 457 | 0.9037 | 0.0570 | 0.0329 |
| 7 | 154 | 0.50 | 199 | 0.59 | 104 | 0.57 | 457 | 0.5546 | 0.0461 | 0.0266 |
| 9 | 152 | 0.41 | 177 | 0.53 | 122 | 0.36 | 451 | 0.4313 | 0.0848 | 0.0490 |
| 11 | 185 | 0.37 | 178 | 0.30 | 110 | 0.24 | 473 | 0.3042 | 0.0683 | 0.0394 |
| ***ftn-1(ok485); ets-4(rrr16)* CTRL** | | | | | | | | | | |
| Days at 4°C | Total | Survival | Total | Survival | Total | Survival | Total animals | Average Survival | St. Dev. (+/-) | SEM (+/-) |
| 5 | 119 | 0.99 | 107 | 1.00 | 120 | 0.98 | 346 | 0.9889 | 0.0127 | 0.0073 |
| 7 | 93 | 0.69 | 105 | 0.63 | 106 | 0.96 | 304 | 0.7597 | 0.1780 | 0.1027 |
| 9 | 110 | 0.50 | 122 | 0.08 | 124 | 0.08 | 356 | 0.2209 | 0.2417 | 0.1396 |
| 11 | 88 | 0.02 | 97 | 0.03 | 138 | 0.08 | 323 | 0.0445 | 0.0308 | 0.0178 |
| ***ftn-1(ok485); ets-4(rrr16)* 30 mM FAC** | | | | | | | | | | |
| Days at 4°C | Total | Survival | Total | Survival | Total | Survival | Total animals | Average Survival | St. Dev. (+/-) | SEM (+/-) |
| 5 | 140 | 0.34 | 73 | 0.49 | 97 | 0.77 | 310 | 0.5364 | 0.2184 | 0.1261 |
| 7 | 144 | 0.00 | 124 | 0.00 | 127 | 0.21 | 395 | 0.0709 | 0.1227 | 0.0709 |
| 9 | 117 | 0.00 | 119 | 0.00 | 74 | 0.00 | 310 | 0.0000 | 0.0000 | 0.0000 |
| 11 | 102 | 0.00 | 116 | 0.01 | 88 | 0.13 | 306 | 0.0445 | 0.0698 | 0.0403 |

Wilcoxon signed rank test

| | *ets-4(rrr16)* CTRL | *ets-4(rrr16)* 30 mM FAC | *ftn-1(ok485); ets-4(rrr16)* CTRL | *ftn-1(ok485); ets-4(rrr16)* 30 mM FAC | wt CRTL |
|---|---|---|---|---|---|
| *ets-4(rrr16)* 30 mM FAC | 0.00049 | | | | |
| *ftn-1(ok485); ets-4(rrr16)* CTRL | 0.00384 | 0.67725 | | | |
| *ftn-1(ok485); ets-4(rrr16)* 30 mM FAC | 0.00252 | 0.00249 | 0.00534 | | |
| wt CRTL | 0.00592 | 0.16817 | 0.01108 | 0.02831 | |
| wt 30 mM FAC | 0.00251 | 0.00049 | 0.00252 | 0.81206 | 0.01507 |
